# Pin4 Links Post-transcriptional and Transcriptional Responses to Energy Depletion in Yeast

**DOI:** 10.1101/2024.11.13.623376

**Authors:** Michaela Ristová, Katherine Bexley, Johanna Franziska Seidler, Vadim Shchepachev, Sushma Sharma, Andrei Chabes, Atlanta G. Cook, David Tollervey

## Abstract

Adaptation to environmental change is essential in all organisms, with RNA-binding proteins (RBPs) playing critical roles in rapid cellular responses. We analyzed the largely uncharacterized yeast RBP Pin4, and its involvement in adaptation to glucose depletion. UV crosslinking (reCRAC) revealed that in glucose conditions Pin4 binds mRNA 3’ UTRs with preference for a specific motif in mRNAs involved in glycolysis, amino acid, and mitochondrial metabolism. Following glucose withdrawal, Pin4-RNA binding was greatly reduced, with residual binding favoring transcripts associated with protein translation. Cells lacking Pin4 were greatly impaired in recovery from nutrient starvation and hypersensitive to oxidative stress, consistent with the mRNA targets. RNAseq and reporter assays indicated that loss of Pin4 correlated with some increases in target mRNA abundance. In wildtype yeast, glucose depletion induces diauxic shift, with massive changes in transcription patterns. Unexpectedly, this response was almost entirely abolished in cells lacking Pin4, its C-terminal prion-like domain, the RNA-recognition motif (RRM) or with an RRM point mutation. Pin4 is implicated in sensing energy depletion, which also occurs during the approach to stationary phase. We postulate that Pin4 helps coordinate post-transcriptional and transcriptional responses to energy stress, via riboregulation.

## INTRODUCTION

The capacity to adapt to ever-changing environments is a fundamental trait necessary for the survival of all living organisms. This adaptability encompasses specific responses to a wide range of stressors such as nutrient depletion, temperature fluctuations, and pH changes (Fulda et al. 2010). Central to these adaptive responses are RNA-binding proteins (RBPs), which regulate a multitude of critical steps throughout gene expression (Gebauer et al. 2020; Kilchert et al. 2020; Birot et al. 2021; Goswami et al. 2024).

The budding yeast *Saccharomyces cerevisiae* grows on plant surfaces and is exposed to many environmental insults. For example, a rain shower will result in immediate nutrient deprivation, perhaps only transiently. Yeast has long been used as a model eukaryote, to study responses to a multitude of stressors. Notable among these stress responses is the diauxic shift, the metabolic switch induced by glucose depletion, which drastically affects yeast cell physiology and behavior (Galdieri et al. 2010). Under glucose-rich conditions, yeast cells metabolize glucose in preference to any other available carbon sources, utilizing aerobic fermentation. Glycolysis converts the glucose to CO_2_ and ethanol, which are excreted despite the availability of oxygen. Metabolic preference for aerobic fermentation – termed the Crabtree effect – has been observed across diverse biological systems, including plant pollen, Trypanosomatids, skeletal muscles, and tumor cells, where it is referred to as the Warburg effect (Cazzulo 1992; Tadege and Kuhlemeier 1997; Alfarouk et al. 2014; Liberti and Locasale 2016). The predominant hypothesis for the emergence of the Crabtree effect centers on the rapid production of ATP (Pfeiffer and Morley 2014). Following glucose depletion, the switch from fermentation to respiration involves a two-stage response: a rapid post-transcriptional response, followed by a slower transcriptional response. The initial post-transcriptional response primarily affects protein synthesis, followed by alterations in mRNA splicing, polyadenylation, RNA stability, RNA modifications, spatial localization, and condensate formation (Ashe et al. 2000; Kuhn et al. 2001; Chang and Huh 2018; Birot et al. 2021; Hernández-Elvira and Sunnerhagen 2022; Heinrich et al. 2024). The subsequent transcriptional response unfolds over several minutes, with alterations in expression of a large set of genes (DeRisi et al. 1997). Key changes include upregulation of specific stress-response genes and suppression of genes involved in ribosome biogenesis and growth.

Central to the regulation of this response are three major signaling pathways associated with yeast glucose metabolism: the Snf1, PKA, and Snf3/Rgt2 pathways (Rolland et al. 2002; Kim et al. 2013; Conrad et al. 2014; Kayikci and Nielsen 2015).

Supporting RNA binding proteins as central to rapid adaptive responses, a recent study identified extensive remodeling of the yeast RNA-protein interactome in response to glucose withdrawal or heat shock within a 16 min timeframe (Bresson et al. 2020). Key factors in protein synthesis were notably affected, consistent with the widespread shutdown of translation during stress. Intriguingly, among the most strongly altered RBPs, several were largely uncharacterized, notably Pin4, Mrn1, Pbp2 and Nab6. Subsequent investigations implicated Mrn1 and Nab6 in a novel post-transcriptional cell wall integrity pathway (Bresson et al. 2023). Here, we characterize the role of Pin4 in the cellular response to glucose withdrawal.

Previous screens identified Pin4 as a protein that alters yeast prion formation and as a Rad53 interactor. Furthermore, Pin4 is implicated in the responses to methylmethane sulfonate (MMS) and bleomycin treatment, and normal G2/M cell cycle progression (Derkatch et al. 2001; Pike et al. 2004; Pike and Heierhorst 2007). However, these putative functions were not clearly related to RNA interactions, so we sought to investigate these in detail. We conclude that Pin4 has a pivotal role in yeast energy metabolism, regulating both post-transcriptional and transcriptional adaptations to cellular stress induced by glucose depletion. Identifying the function of this critical protein highlights the potential for a class of previously unidentified RNA binding proteins that connect external signaling to a regulatory RNA interaction network.

## RESULTS

### Pin4 has a non-canonical RRM and large unstructured regions

Since very little functional information was available for Pin4, we first sought structural features that might point to its role in the cell. Conserved domains were analyzed utilizing multiple sequence alignments from the NCBI Conserved Domain Database (Wang et al. 2022). Pin4 is 668 amino acids in length, with an unstructured Q/N-rich C-terminal region previously identified as a prion-forming domain (Derkatch et al. 2001; Yang et al. 2013, 2014). Database searches and AlphaFold3 predicted three further domains: an RNA Recognition Motif (RRM), a coiled-coil (CC) domain, and an R3H domain with all other parts being largely unstructured (Fig. 1A) (Wang et al. 2022; Abramson et al. 2024).

**Figure 1:**
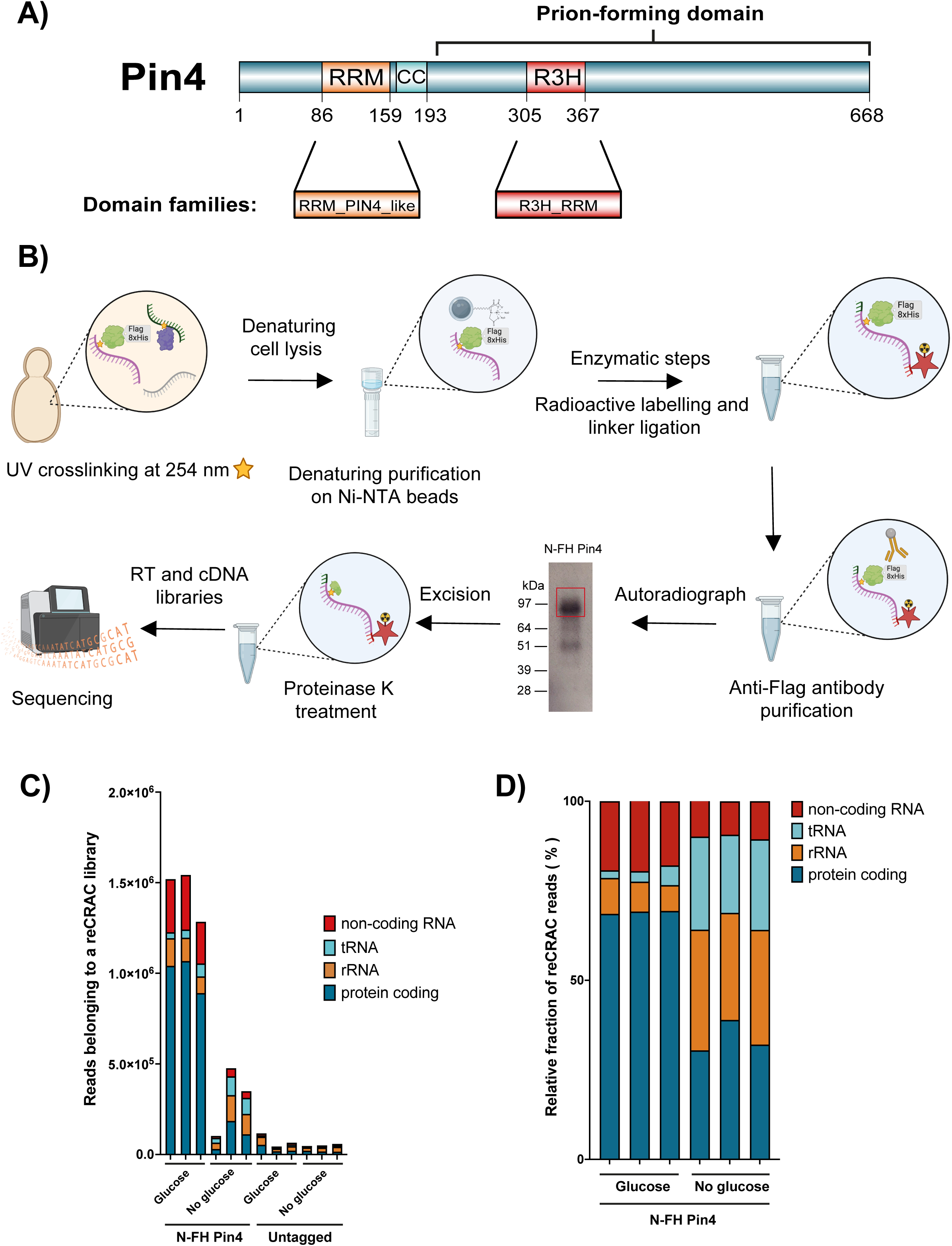
Identification of the Pin4 RNA-bound interactome A) Pin4 is a largely unstructured protein with three predicted domains, an RNA Recognition Motif (RRM), a coiled-coil domain (CC), and an R3H domain. The previously identified prion- forming domain is also annotated. The RRM and R3H domain regions belong to protein domain families, RRM_PIN4_like and R3H_RRM respectively. B) Overview of the reCRAC technique. Briefly, yeast cells were irradiated at 254 nm to induce covalent crosslinks (yellow stars) between proteins and associated RNAs. Cells were lysed in denaturing conditions, to prevent protein degradation and reduce non-specific interactions. The bait protein carries a FLAG-His8 tag, allowing purification on Ni-NTA beads, where the enzymatic steps of RNA digestion, radiolabeling and linker ligation (red star) were performed. This was followed by elution and purification on anti-FLAG beads. Complexes were gel-purified, the radioactive RNA visualized by autoradiography and the region corresponding to the RNA-protein complexes excised. The bait protein was treated with Proteinase K, leaving one or more residual aminoacids. RNA was extracted, converted to cDNA by reverse transcription (RT), and sequenced. See the methods section for more detail (Ristová et al. 2025). Figure created with BioRender. C) Total reads recovered in reCRAC libraries from Glucose and No glucose conditions (16 min in Gly/EtOH) for FH-Pin4 and Untagged control. Each condition was analyzed in triplicate. The Untagged background control had minimal number of reads as expected. D) The relative fraction of reCRAC reads belonging to Glucose and No glucose (16 min in Gly/EtOH) Pin4 libraries. The experiment was performed in triplicate for each condition.

The Pin4 RRM is from the family “RRM_PIN4_like” domain (Fig. 1A) (Wang et al. 2022), which is also present in five annotated proteins from fungal species and one protozoan (Supp. Figs. 1A and 1B). Of these, only *Schizosaccharomyces pombe* Cip1 and Cip2 are characterized, and were reported to function in gene regulation during oxidative stress (Martín et al. 2006). Interestingly, the Pin4 RRM differs from canonical RRM domains, with alterations in the characteristic ribonucleoprotein (RNP) motifs. Positions 3 and 5 in RNP-1 and position 2 in RNP-2 are typically aromatic residues, Phe or Tyr, important for RNA binding. In Pin4, the corresponding amino acids are Leu128 (RNP-1 position 3), Phe130 (RNP-1 position 5), and Val 88 (RPN-2 position 2) (Supp. Figs. 2A and 2B). Although, the leucine and valine residues are not aromatic, but hydrophobic, it is not completely unusual as equivalent residues can be also found in the same positions in RRMs of other yeast RNA binding proteins, including UPF3 and TAP (Maris et al. 2005).

The Pin4 R3H is from the family of “R3H_RRM” domain, identified in other ten fungal proteins, mostly uncharacterized except for *S*.*pombe* Cip2 (Supp. Figs. 1A and 1B). R3H domains can dimerize (Liu et al. 2007; He et al. 2013) and AlphaFold3 reproducibly modeled the R3H domain of Pin4 as a homodimer, with good supporting statistics (Supp. Fig. 2C).

This suggests that the R3H domain of Pin4 may be a dimerization element rather than a nucleic acid binding domain. The CC domain is predicted to mediate protein interactions, and also lies close to the RRM (Supp. Fig. 2D) (Heierhorst 2008), indicating a scaffolding role that could facilitate spatial proximity between RNA targets and protein interactors.

### Pin4 predominantly binds mRNA 3’ untranslated regions

Pin4 was an outlier in the total RNA-bound proteome following stress (Bresson et al. 2020). We therefore identified RNAs bound by Pin4 during growth in 2% glucose medium and following transfer to 2% glycerol plus ethanol (Gly/EtOH) medium for 16 min, to induce diauxic shift. To allow purification, we expressed Pin4 as a fusion protein with an N-terminal tandem-affinity tag containing a single FLAG epitope-Ala_4_-His_8_ (FH tag) (Bresson et al. 2020). The 20 amino acid FH tag was introduced into the chromosomal *PIN4* gene under control of the endogenous promoter, using CRISPR-Cas9 without a selective marker (Laughery et al. 2015). FH-Pin4 is the only form of Pin4 in these cells, which showed no clear growth defects.

We initially applied the published protocol for crosslinking and analysis of cDNA (CRAC) (Granneman et al. 2009). However, Pin4 underwent substantial degradation during purification (Supp. Fig. 3A). We therefore developed a modified protocol, termed reverse CRAC (reCRAC) (Ristová et al. 2025), which notably enhanced the integrity of Pin4 (Fig. 1B, Supp. Fig. 3A). In reCRAC, the tandem affinity purifications steps are reversed relative to CRAC. Briefly, cells are lysed in highly denaturing buffer, including 6M Guanidine-HCl, to immediately inactivate cellular proteases. FH-Pin4 was recovered on Ni-NTA beads in denaturing buffer. While still immobilized on the Ni-NTA beads, but under non-denaturing conditions, bound RNAs were subjected to partial Benzonase digestion, the 5’ ends were labeled with ^[32]^P, and linkers were added to the termini. Protein-RNA complexes were eluted and further purified on an anti-FLAG column. Following elution, complexes were gel purified using the labeled RNA as marker, and the proteins were degraded with proteinase K. RNA was recovered and used for RT-PCR amplification and sequencing. Replicate experiments showed good reproducibility (Supp. Figs. 3B and 3C).

During reCRAC, samples were barcoded and mixed prior to RT-PCR amplification. In consequence, relative numbers of total reads recovered for each condition approximates to overall crosslinking efficiency. This allows direct comparison of samples. The total number of reads in the FH-Pin4 libraries decreased dramatically following glucose withdrawal, indicating strongly decreased overall RNA binding (Fig. 1C). In glucose rich conditions, Pin4 predominantly targeted mRNAs, which constituted about 70% of its interactions (Fig. 1D).

Reads were also mapped to “non-coding” RNAs, but these were distributed over a large and heterogeneous set of RNAs, mainly antisense and intergenic, and specific interactions were not evident. Following glucose depletion, there was a 50% reduction in the fraction of reads mapped to mRNA (Fig. 1D). Absolute recovery of rRNA was similar to the untagged control, indicating that these represent non-specific interactions (Supp. Fig. 3D), and the apparent increase following glucose depletion likely reflects loss of mRNA binding. Pin4 protein levels remained constant during the experiment in both glucose and glucose starvation conditions (Supp. Fig. 3E).

To assess Pin4 binding specificity, we analyzed the distribution of FH-Pin4 bound sequencing reads across the 5’ untranslated region (UTR), coding sequence (CDS), and 3’ UTR of all annotated mRNAs. These were compared to poly(A)^+^ selected mRNA sequencing (RNAseq) obtained under the same conditions. At short time points following glucose withdrawal, abundances of most mRNAs were little altered, as previously reported; stress response RNAs accumulate substantially only at later time points and previously synthesized mRNAs are stabilized, probably through sequestration into condensates (Zid and O’Shea 2014; Bresson et al. 2017, 2020; Zedan et al. 2024; Glauninger et al. 2024).

In the presence of glucose, Pin4 predominantly bound 3’ UTR regions, consistent with a previous report (Hogan et al. 2008); 3’ UTRs formed 78% of reCRAC reads compared to 12% in total RNA from the RNAseq control (Fig. 2A). This pattern of predominant 3’ UTR binding is apparent in a metagene plot in which reCRAC reads are mapped onto mRNAs aligned at their 3’ ends, revealing a distinct 3’ peak (Fig. 2B). On individual mRNAs with strong Pin4 binding relative to RNAseq, the prominent peak in the 3’ UTR is very clearly visible (shown for *TPI1*, *ILV5*, *GPM1* and *RPS12* in Fig. 2C).

**Figure 2:**
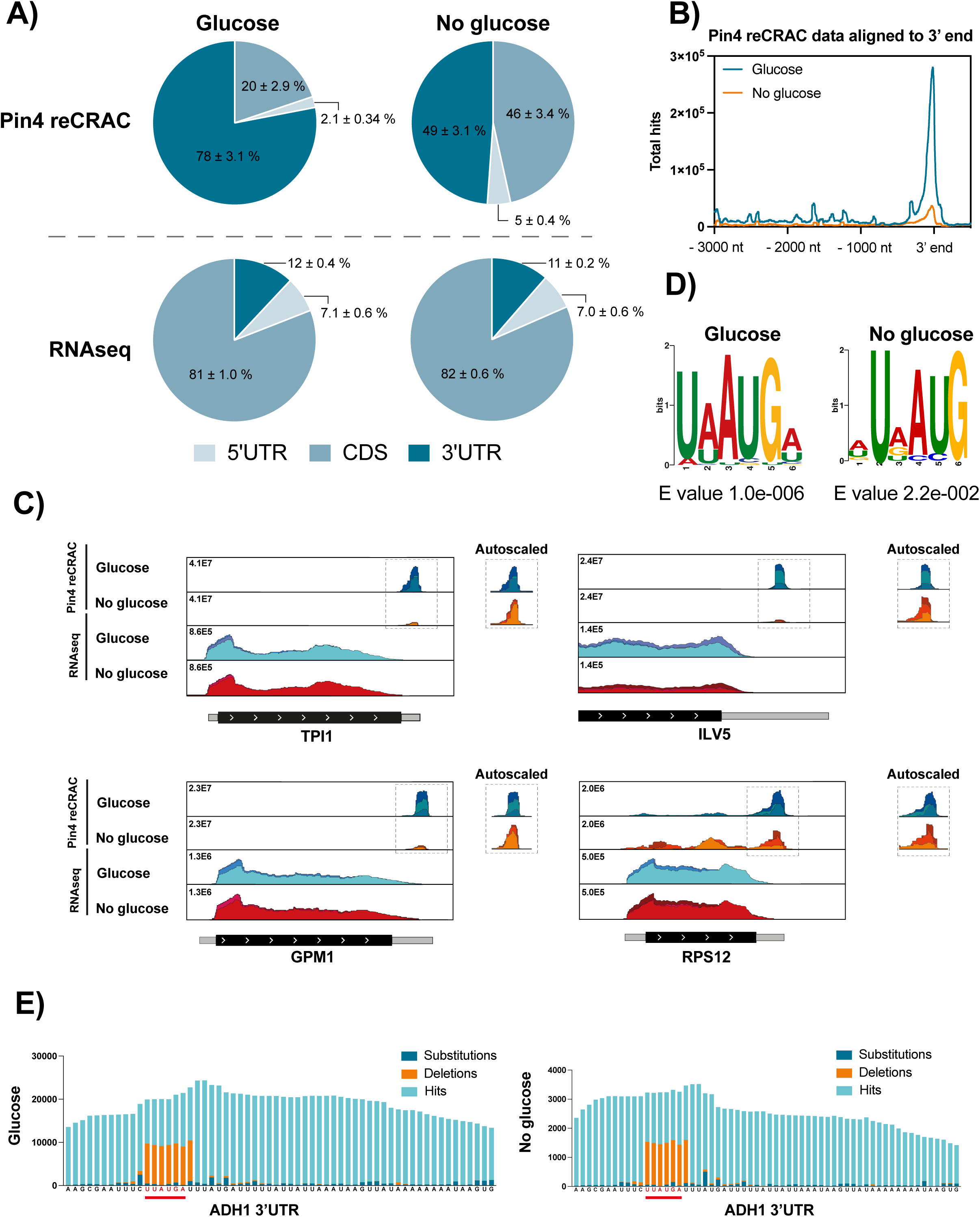
Pin4 predominantly targets a single motif in 3’ UTRs A) Distribution of Pin4 binding to 5’UTR, CDS, and 3’ UTR of target mRNAs in Glucose and No glucose (16 min in Gly/EtOH) conditions. RNAseq was included as a control. Pin4 targets mainly 3’ UTRs in both conditions. B) Metagene plot showing the total hits across Pin4 reCRAC data aligned to the 3’ end in Glucose (blue) and No glucose (orange) conditions. The highest peak is present close to the 3’ end in both conditions, but substantially lower following glucose depletion. C) Examples of FH-Pin4 binding across its top targets (*TPI1*, *ILV5*, *GPM1*, *RPS12*). Three replicates are plotted on top of each other, each shown in a different color shade. RNA-seq is shown for comparison. The reads are normalized to the size of the library and scaled to the highest peak in Pin4 reCRAC and RNAseq. The autoscaled Pin4 reCRAC peaks are shown on right next to the plots. D) Pin4 motif (MEME) found in the 3’ UTR of Pin4 targets (mRNA species that individually comprise ≥ 0.3% of the total library) in Glucose (35 RNAs) and No glucose (38 RNAs) conditions. The E value is given below each motif. E) Close up view of reCRAC pileups of the 3’ UTR of Pin4 target, *ADH1*, in Glucose and No glucose conditions. The motif sequence UA/UAUGA/U is marked in red and underlined. The deletion peaks (orange) represent the sites of crosslinks between RNA and protein.

Following glucose withdrawal, the enrichment for Pin4 binding to 3’ UTRs is notably decreased (49% of all reads) (Fig. 2A), although some indication of preferential 3’ binding remained (Fig. 2B). On most strongly bound mRNAs, residual Pin4 association remained enriched on the 3’ UTR (Fig. 2C). However, a subset had substantial CDS binding after glucose depletion. These transcripts were generally related to ribosomes and translation (Supp. Fig. 3F; Supp. Table 4). As an example, *RPS12* (Fig. 2C) has substantial Pin4 binding in the CDS, even though the highest signal was still in the 3’ UTR.

### A binding motif for Pin4

In reCRAC, the direct sites of RNA-protein crosslinks can be pinpointed by searching for deletions within recovered sequences. At the end of the experimental procedure, the bound protein is degraded by proteinase K treatment, leaving at least one amino acid on the RNA. Due to this, reverse transcriptase can skip the crosslinked nucleotide. Nucleotide substitutions can arise from other causes (PCR, RT, sequencing) but single nucleotide deletions are characteristic, and otherwise very rare. Using this data on Pin4 binding sites to identify potential sequence determinants, we focused on strong Pin4 targets under both glucose and Gly/EtOH conditions; the cutoff for motif analysis being at least 0.3% of the library in at least two replicates (35 transcripts in glucose; 38 transcripts following withdrawal) (Supp. Table 5). After identifying the most significant deletion peaks within these transcripts, we ran MEME analysis on the flanking 20 nucleotide sequences. We discovered a consistent U(A/U)AUG(A/U) motif across both conditions (Fig. 2D). This motif was generally located either precisely at the peak of binding (Fig. 2E), or immediately adjacent to the highest deletion peak (Supp. Fig. 3G). The close association of the motif with the peak of Pin4 binding was maintained in glucose replete and depleted conditions. Analysis of mRNAs that were well bound by Pin4, but not enriched relative to mRNA abundance (on or below the diagonal in Figs. 3A and 3B), showed no clear motif enrichment.

**Figure 3:**
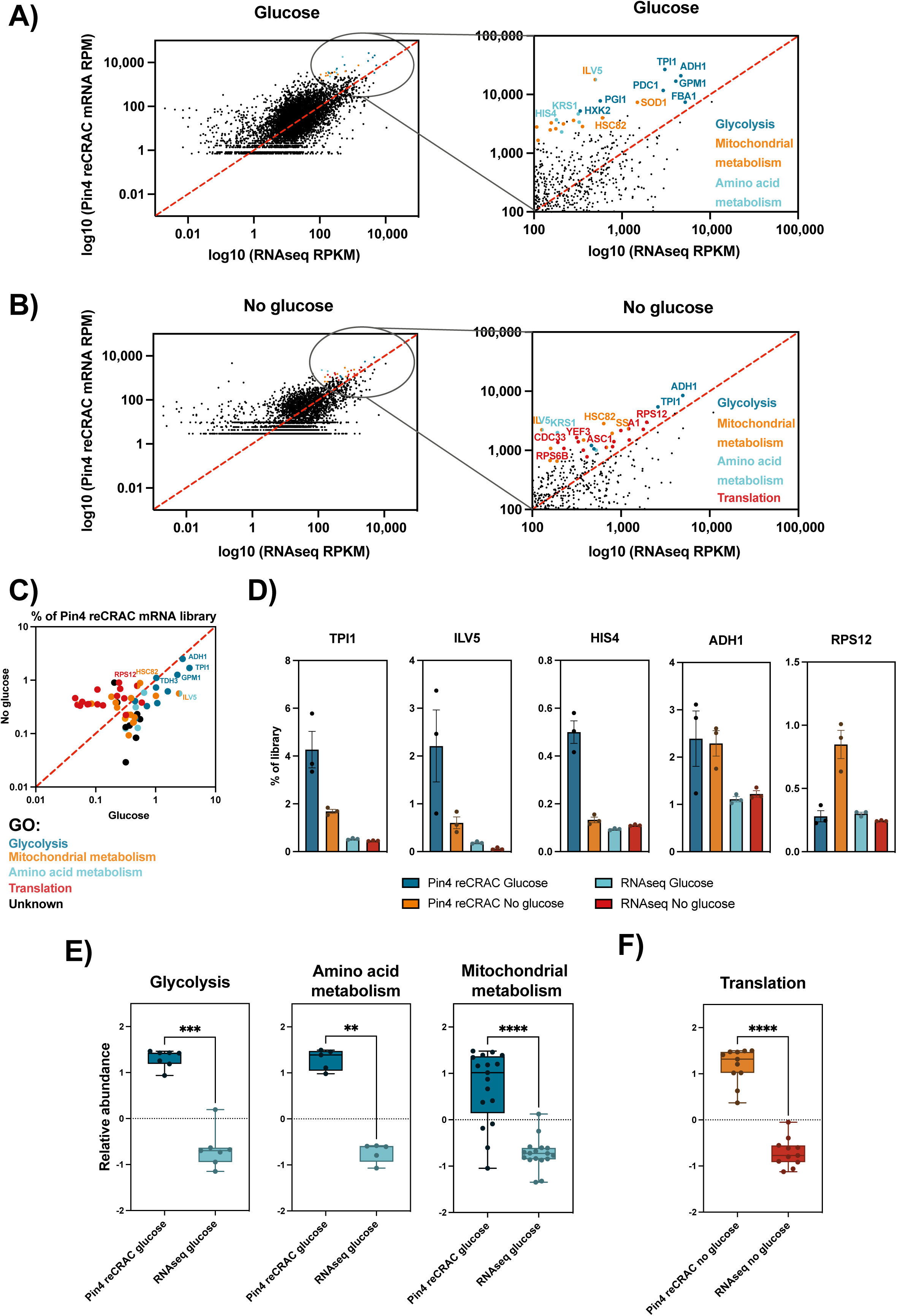
Pin4 binding is enriched on metabolic and growth-associated mRNAs A-B) Scatterplots comparing the changes in Pin4-RNA binding versus the mRNA abundance in Glucose and No glucose (16 min in Gly/EtOH) conditions. The zoomed in window shows the top Pin4-RNA enriched targets colored based on their GO term enrichment. C) Scatterplot showing the top mRNA Pin4 targets (≥ 0.3% of the total library) in glucose and glucose depletion (16 min) conditions. The transcripts fall into four GO term enrichment categories: glycolysis, mitochondrial and amino acid metabolism, and translation. D) Bar graphs showing the percentage of the total Pin4 reCRAC library, glucose or glucose depleted samples, together with the RNAseq control. Mean and individual replicates are plotted. Error bars show SEM (n=3). E-F) mRNAs with good recovery in Pin4 reCRAC reCRAC (≥ 0.3% of the total library and ≥1.5-fold enrichment in ≥2 replicates) were subjected to GO term analysis. The box plots show scaled abundance in reCRAC relative to RNAseq, for mRNAs identified under different GO terms. Significant differences between conditions were calculated with the Mann-Whitney statistical test and are indicated by asterisks (** p < 0.01, *** p < 0.001, **** p< 0.0001). Individual replicate values are plotted.

Analysis of the strong Pin4 targets showed no consistent pattern of pairs of binding sites that would indicate binding as a homodimer. Moreover, on mRNA targets where we could confidently map the precise crosslinking position, this was over a single motif (Fig. 2E and Supp. Fig. 3G). We conclude that the U(A/U)AUG(A/U) motif is very likely to represent a direct Pin4 recognition element.

### Pin4 selectively binds metabolic and growth-associated transcripts

To assess mRNA target specificity, Pin4-associated RNAs isolated with reCRAC were compared to mRNA abundances from WT RNAseq (Supp. Table 6; Figs. 3A and 3B). The majority of mRNAs were detectably associated with Pin4. The most enriched Pin4- associated transcripts in glucose medium were predominantly involved in metabolic and pro-growth pathways (Fig. 3A). Strongly bound targets (≥0.3% of library) that showed enrichment in reCRAC relative total mRNA (≥1.5-fold in ≥2 replicates) are highlighted. Key glycolytic enzymes emerged as top targets in both conditions, including *TPI1* (triose phosphate isomerase), *ADH1* (alcohol dehydrogenase), and *GPM1* (phosphoglycerate mutase) (Fig. 3C). *TPI1* alone represented about 4% of reCRAC reads, versus 0.5% from RNAseq in glucose (Fig. 3D). Genes involved in amino acid metabolism (e.g., *HIS4*) were strongly targeted only in glucose-rich conditions, as were nuclear-encoded transcripts involved in mitochondrial function (e.g., *ILV5*). However, no mitochondrial encoded RNAs were well recovered.

For strong interactions (≥0.3% of library and ≥1.5-fold enrichment in reCRAC over RNAseq in ≥2 replicates) (Supp. Table 5) comparing Pin4 binding relative to transcript abundance, confirmed significant GO term enrichment: glycolysis (p=0.0006), amino acid metabolism (p=0.0079), and mitochondrial metabolism (p<0.0001) (Fig. 3E). This enrichment was also reproduced by grouping the transcripts based on their binding enrichment and GO term analysis using the Proteomaps tool (Liebermeister et al. 2014) (Supp. Fig. 4A). These strong targets showed good reproducibility across replicates in both conditions (Supp. Fig. 4B).

Following glucose withdrawal, absolute binding was reduced on almost all mRNAs, and there were also fewer relatively enriched targets, consistent with the metagene plot. However, some of the same genes remained relatively enriched, including the glycolytic enzymes. Representation was strongly increased for several translation-related transcripts (≥0.3% of library and ≥1.5-fold enrichment in ≥2 replicates) (Fig. 3F); these correspond to the mRNAs showing Pin4 association across the CDS (e.g. *RPS12*) (Fig. 2C).

### Pin4-deficient cells are sensitive to glucose depletion and mitochondrial stress

As Pin4 binds mRNAs connected to central carbon metabolism and mitochondria, we tested for growth phenotypes after precisely deleting *PIN4* using a KanMX6 cassette (Bähler et al. 1998). In glucose medium, *pin4Δ* strains showed only a slight increase in doubling time compared to WT (Supp. Fig. 5A). However, upon shifting to 2% Gly/EtOH medium, *pin4Δ* cells exhibited greatly increased lag before reinitiation of growth (Supp Fig. 5B), followed by a steady-state doubling time of approximately 7 hr versus 6 hr for the WT (Supp. Fig. 5C).

Under oxidative stress induced by 0.5 mM hydrogen peroxide, *pin4Δ* cells had a lag phase of 30 hr compared to 15 hr for WT (Supp. Figs. 5D and 5E). At 1 mM H_2_O_2_, *pin4Δ* cells showed no growth, whereas WT cells remained viable. Exposure to glyoxal, which collapses mitochondrial membrane potential, also led to slowed growth and increased lag phase in *pin4Δ* cells (Supp. Fig. 5F), with growth progressively decreasing at higher concentrations compared to WT (Supp. Fig 5G).

We conclude that cells lacking Pin4 exhibit hypersensitivity to glucose withdrawal or mitochondrial stress, consistent with the targets for Pin4 binding.

### Pin4 negatively regulates target mRNA abundance

To determine whether the loss of Pin4 affects target mRNA abundance, we performed RNA sequencing on WT and *pin4Δ* cells in glucose medium. Differential expression (DE) analysis identified 58 mRNAs showing the most altered expression between the WT and *pin4Δ* (fold change ≥ 2.0; adj. p ≤ 0.05) (Fig. 4A; Supp. Table 7). The most upregulated mRNAs were enriched for cell wall components (Supp. Fig. 6A), but these generally had very low expression, making the significance unclear. Downregulated genes were preferentially linked to transmembrane transport and vitamin binding (Supp. Fig. 6B). The mRNA showing strongest DE were not top Pin4 targets. However, comparison of Pin4 mRNA binding and changes in abundance in *pin4Δ* cells revealed generally increased abundance for preferential Pin4 targets (strong mRNA targets with at least 0.3% of the library) (Fig. 4B).

**Figure 4:**
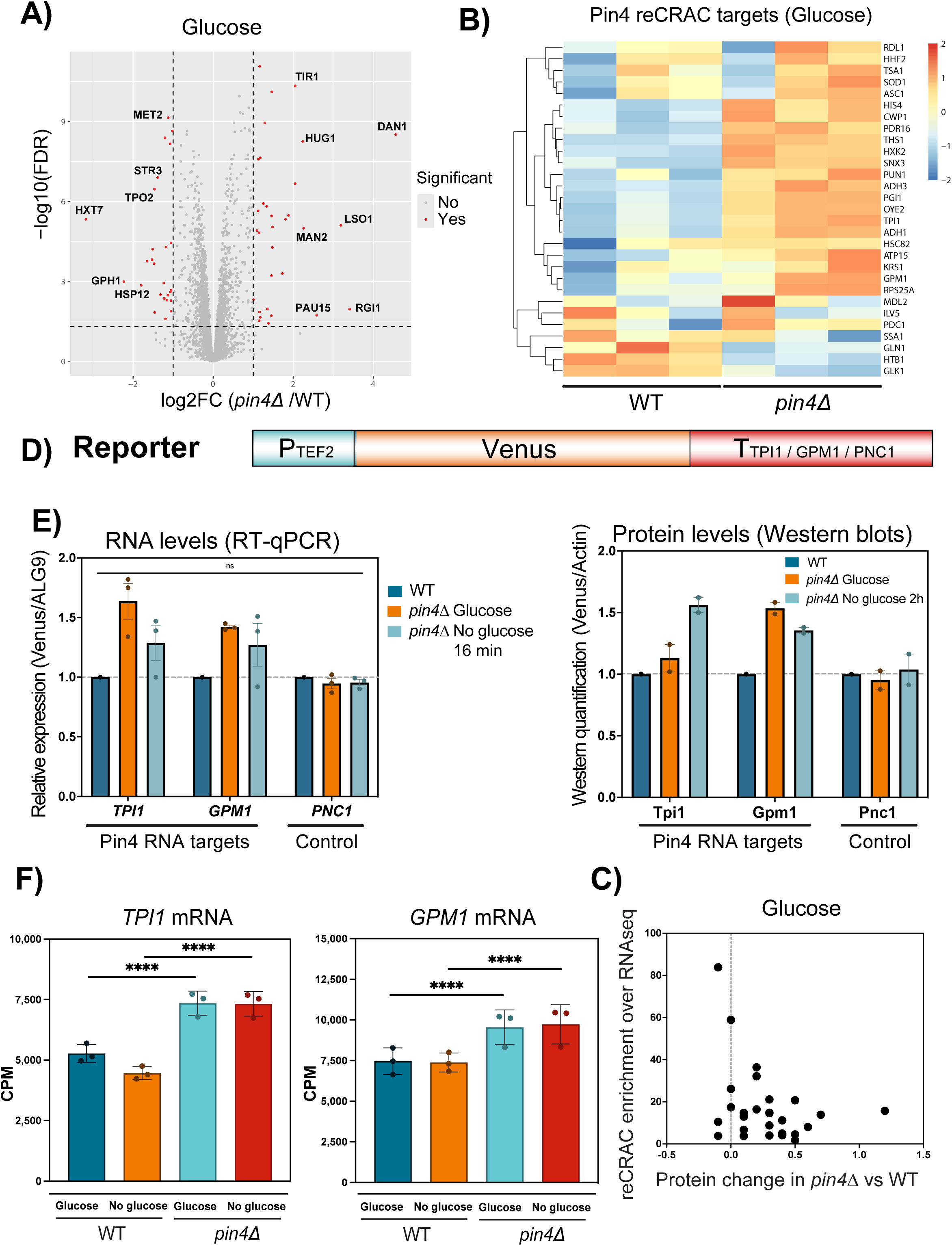
Pin4 negatively regulates the abundance of target mRNAs A) Volcano plot of differential gene expression analysis in *pin4Δ* compared to the WT under glucose conditions. Significantly downregulated and upregulated genes are highlighted in red (adj. p value ≤ 0.05, log2FC ≥ 2). B) Heatmap showing the expression of top Pin4-RNA (≥ 0.3% of the total library and ≥1.5-fold enrichment in ≥2 replicates) targets in WT and *pin4Δ* under glucose condition. Each row represents a different gene and each column represents a different experimental replicate. Expression levels are color-coded from blue (low expression) to red (high expression) and normalized. C) Scatterplot showing the relationship between Pin4 RNA binding and protein abundance for top target mRNAs during growth in glucose. The x-axis represents the log2 fold change in protein abundance in *pin4Δ* cells (from published proteomics data), while the y-axis shows fold Pin4 binding enrichment (reCRAC normalized over RNAseq) for transcripts representing ≥0.3% of the reCRAC library and ≥1.5-fold enrichment. D) Schematic representation of the reporter construct used in this study. The construct contains the P_TEF2_ promoter, driving the expression of the Venus fluorescent protein, which is fused to the 3’ UTR from top Pin4-RNA targets: *TPI1* and *GPM1*. As control, the *PNC1* terminator sequence was used. E) Bar graphs showing the levels of RNA (left) and protein (right) for specific genes (*TPI1* and *GPM1*) that are targets of Pin4 regulation, alongside a control (*PNC1*). Data are represented for WT and *pin4Δ* strains under glucose and glucose depletion conditions (16 min for RNA levels (n=3), 2 hr for protein levels (n=2)). Values are normalized to the WT in glucose (dark blue). Significance was calculated using Mann Whitney two-tailed test. Individual replicate values are plotted. F) Bar graphs depicting the mRNA levels of *TPI1* and *GPM1* in WT and *pin4Δ* in glucose and glucose depletion conditions as determined from RNAseq and normalized by counts per million (CPM) (n=3, mean values and replicates plotted; error bars represent SD). Each dot represents an individual replicate. Statistical analyses were performed using EdgeR and significant differences were observed with FDR-adjusted p-values of 2.60 × 10⁻^8^ and 1.98 × 10⁻^11^ for TPI1 in glucose and glucose depletion between WT and *pin4Δ* strains, respectively, and 2.43 × 10⁻^7^ and 3.31 × 10⁻^9^ for GPM1. Mean and individual replicates are plotted.

**Figure 5:**
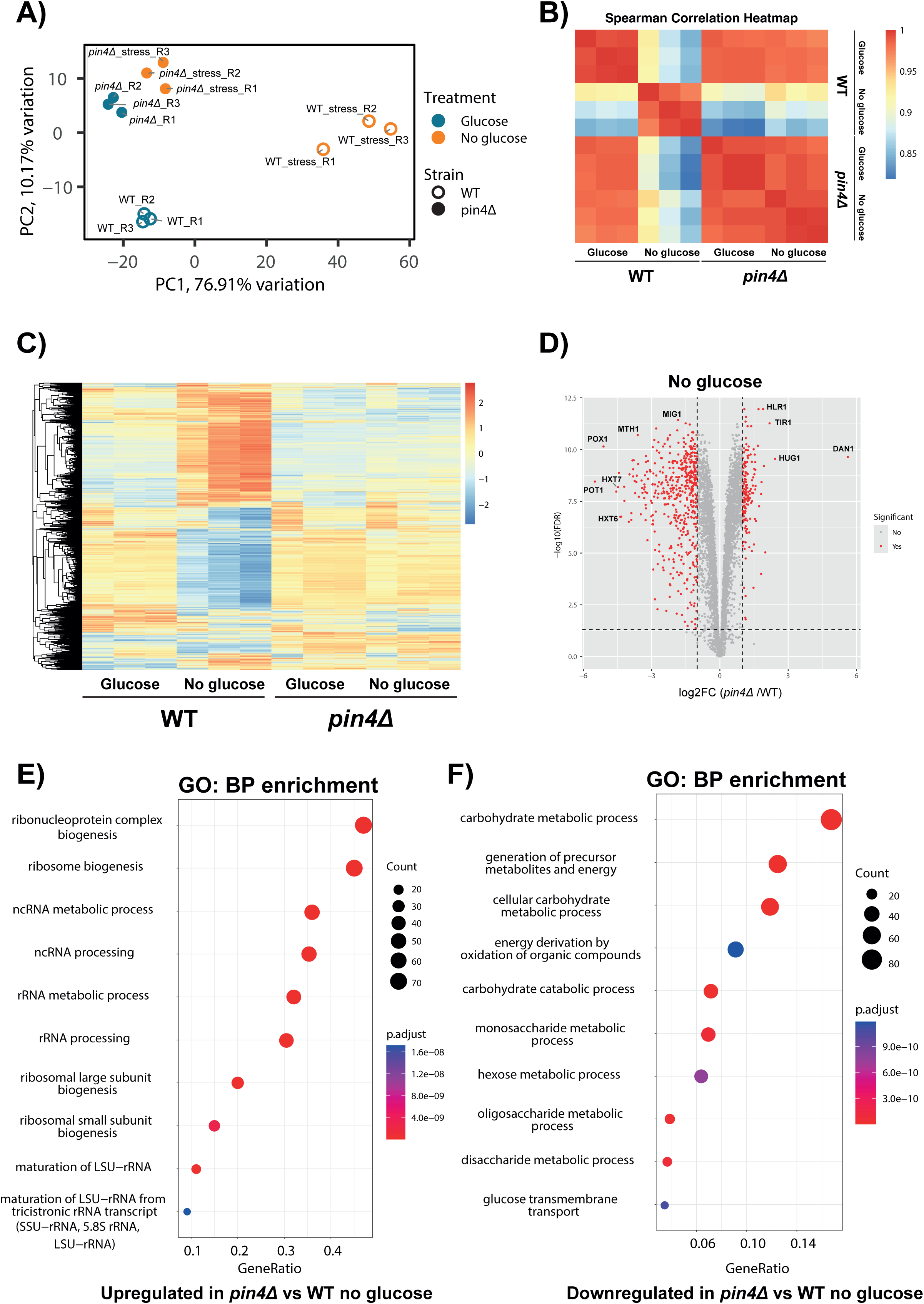
Loss of Pin4 blocks transcriptional responses to glucose depletion A) Principal component analysis (PCA) depicting the variation in gene expression profiles between the WT and *pin4Δ* with and without glucose (16 min in Gly/EtOH). Each point represents a biological replicate, showing the distinct clustering of each group based on their overall gene expression. B) Spearman correlation heatmap showing pairwise comparisons of gene expression between WT and *pin4Δ* under different conditions. The heatmap colors represent Spearman correlation coefficients, indicating the degree of similarity between the conditions. C) Heatmap of gene expression across WT and *pin4Δ* strain in glucose and glucose depletion conditions. Genes are clustered by the rows, based on expression patterns. The color scale represents expression levels from low (blue) to high (red). D) Volcano plot of differential gene expression analysis in *pin4Δ* compared to the WT under glucose depletion conditions (16 min in Gly/EtOH). Significantly downregulated and upregulated genes are highlighted in red (adj. p value ≤ 0.05, log2FC ≥ 2). E) Bubble chart representing the GO term enriched biological processes among mRNAs with increased levels in *pin4Δ* relative to the WT. The size of each bubble indicates the gene count, while the color reflects the p-value. Gene ratio is defined as the proportion of differentially expressed genes (DEGs) that are present in a particular gene set, divided by the total number of DEGs. F) As E but showing mRNAs with reduced levels in *pin4Δ*.

Analysis of recent proteomic data for *pin4Δ*, based on a genome-wide deletion screen, also showed increased protein abundances for most products of strong Pin4 mRNA targets (defined as ≥0.3% of library and ≥1.5-fold enrichment over RNAseq under the same conditions in ≥ 2 replicates) (Fig. 4C) (Messner et al. 2023). This suggested that Pin4 binding might destabilize target mRNAs and/or inhibit translation.

To test this hypothesis, we used the MoClo system to generate reporter constructs (Fig. 4D). These express the Venus fluorescent protein under the control of the *TEF2* promoter (P_TEF2_), which is characterized by high and stable expression levels in both glucose and Gly/EtOH media, with expression unaffected in *pin4Δ* cells. We incorporated the 3’ UTR and terminator sequences of two strong Pin4 targets, *TPI1* (T_TPI1_) and *GPM1* (T_GPM1_), extending 479 and 500 nucleotides, respectively, beyond the stop codons. As a control, we used the 3’ UTR and terminator sequence of *PNC1* (T_PNC1_; 379 nucleotides), which is not bound by Pin4. The constructs where chromosomally integrated into the yeast HO locus, and Venus expression was monitored for mRNA (RT-qPCR) and protein (western blot levels).

In strains lacking Pin4, reporter constructs containing *TPI1* and *GPM1* 3’ UTRs showed a mild increase in mRNA levels in both glucose and following transfer to Gly/EtOH (16 min), which was not observed with the *PNC1* control (Fig. 4E). This result was consistent with our RNAseq data (Fig. 4F). Western blot analysis mirrored these findings at the protein level, displaying a modest increase in the expression of reporters with *TPI1* and *GPM1* 3’ UTRs in *pin4Δ* cells in both nutritional states (Fig. 4E; Supp Fig. 6C). Although the increases observed by RT-qPCR failed to pass a rigorous test for statistically significance, the trend of upregulation among Pin4 targets was seen across all analyses; PCR and western on reporters; RNAseq and proteomics on endogenous mRNAs and proteins. Together, these data support the conclusion that Pin4 binding modestly destabilizes associated mRNAs, with consequent reductions in protein abundance.

**Figure 6:**
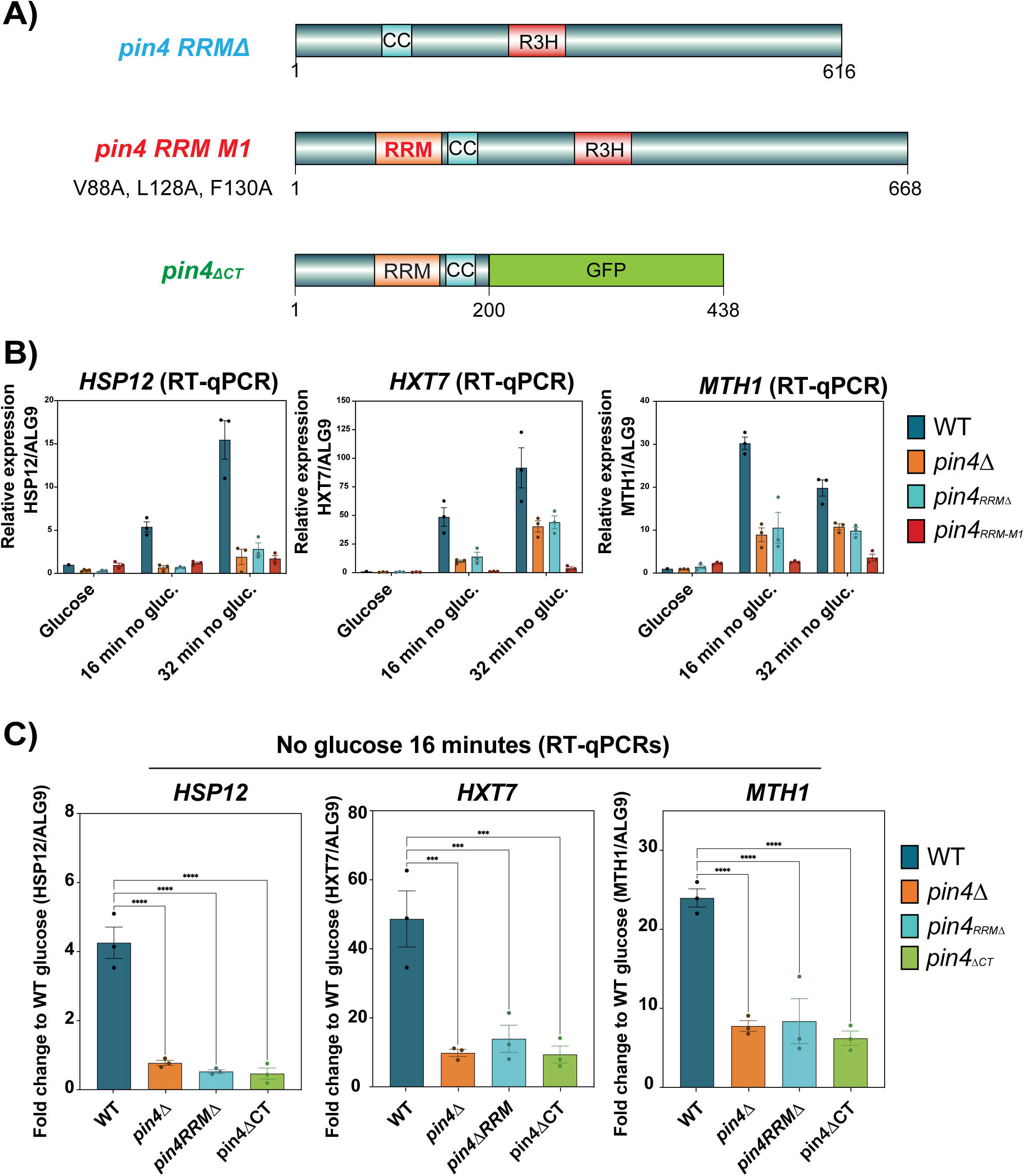
Pin4 RRM and prion-forming domains are critical for the transcriptional response to glucose depletion A) Schematics of representation of constructs, pin4*_RRMΔ_*, lacking the RRM domain, pin4*_RRM-M1_* with mutated RRM domain (in red), and pin4*_ΔCT_*(green). B) Bar graphs comparing RNA expression levels determined by RT-qPCR normalized to *ALG9*, for strains carrying WT *PIN4*, *pin4Δ,* pin4*_RRMΔ_* or pin4*_RRM-M1_* in glucose medium and following glucose depletion for 16 or 32 min (n=3, mean with SEM). Individual replicate values are plotted. C) As B but with strains WT *PIN4*, *pin4Δ,* pin4*_RRMΔ_* or pin4*_ΔCT_*, with glucose depleted for 16 min. Values are plotted as a fold change to WT in glucose conditions (*** p < 0.001, **** p < 0.0001; n=3, mean with SEM). Ordinary one-way ANOVA with Dunnett’s multiple comparisons test was used for the statistical test. Individual replicate values are plotted.

### Impaired transcriptional response to stress in Pin4-deficient cells

Several mRNAs with altered abundance in *pin4Δ* cells were not strongly bound Pin4 targets (Supp Fig. 6D). This indicated that the role of Pin4 involves mechanisms beyond specific interactions with individual mRNA. Wild-type cells undergo profound transcriptional changes following glucose withdrawal, characterized as the integrated stress response (ISR), which alters the abundance of several hundred mRNAs to promote cellular recovery. We analyzed RNA sequencing data following glucose depletion to determine whether the loss of Pin4 affects this response.

Principal component analysis (PCA) revealed surprisingly distinct patterns: WT replicates clustered well in each condition and were clearly separated between glucose replete and glucose depleted conditions. In striking contrast, *pin4Δ* samples clustered together regardless of the treatment (Fig. 5A). This clustering suggested minimal transcriptional changes in *pin4Δ* cells following glucose depletion. Spearman correlation heatmaps confirmed the distinction between WT in glucose versus Gly/EtOH, but high similarities in both conditions for *pin4Δ* cells – as well as consistency in the replicates (Fig. 5B). A heatmap of gene expression (Fig. 5C) further confirmed the minimal variation in *pin4Δ* cells following glucose depletion. This indicates a critical role for Pin4 in the transcriptional response to glucose starvation.

Detailed analysis of all individual transcripts further underscored these observations. Following glucose depletion, differential gene expression (DE) analysis identified 645 transcripts present at significantly different levels in WT and *pin4Δ* cells (fold change ≥ 2.0; adj. p ≤ 0.05), a strikingly large number (Fig. 5D; Supp. Table 8). In almost all cases, DE reflects stress-induced changes, up and down, in the WT that are lost in *pin4Δ*. As expected, stress response genes were strongly upregulated in the WT, whereas pro-growth genes were downregulated, including RNP and ribosome biogenesis. In cells lacking Pin4, the latter were little altered and thus elevated in comparison to the WT (Fig. 5E; Supp. Fig. 7A). Additionally, mRNAs linked to catabolic and respiration-related processes were lower in *pin4Δ* (Fig. 5F; Supp. Fig. 7B).

The duration of the transcriptional defect was tested by RT-qPCR on the *MTH1*, *HSP12*, and *HXT7* genes, which were strongly upregulated in RNAseq data following glucose withdrawal for 16 min in the WT. At 1 hr and 8 hr following glucose withdrawal their expression showed some increase in *pin4Δ*, but remained much lower than in the wildtype (Supp. Figure 8A).

We considered whether *pin4Δ* cells might show greater reliance on oxidative phosphorylation relative to glycolysis during growth on glucose. Wildtype yeast almost exclusively utilizes aerobic fermentation on glucose and is therefore resistant to growth inhibition by Antimycin A (AA), which targets complex III of the mitochondrial electron transport chain. This is also the case for *pin4Δ* (Supp. Fig. 8B), indicating that a metabolic switch does not underlie the differences in stress responses.

### Pin4 domains required for the transcriptional response to glucose depletion

To better characterize the role of Pin4 in the transcriptional response to glucose starvation, we constructed Pin4 mutant strains lacking functional domains.

To assess the role of RNA binding, we created strains precisely lacking the RRM domain (*pin4_RRMΔ_*) or with point mutations (*pin4_RRM-M1_*: V88A, L128A, F130A) predicted to disrupt RNA binding while maintaining RRM domain structure (Fig. 6A). Both constructs were expressed as FH-tagged constructs, and accumulated at elevated levels relative to FH-Pin4 (Supp. Fig. 9A). Unexpectedly, the *pin4_RRM-M1_*variant exhibited notably slower growth than WT, *pin4Δ* or pin4*_RRMΔ_*strains (Supp. Fig. 9B). The basis for this is unclear and currently under investigation.

The effects of these mutations were tested by RT-qPCR on the *MTH1*, *HSP12*, and *HXT7* genes. Expression was analyzed during growth on glucose, and 16 or 32 min after transfer to Gly/EtOH medium. In glucose medium, the mRNAs were expressed at similar, low levels in all strains, WT, *pin4Δ*, *pin4_ΔRRM_* or *pin4_RRM-M1_*. Following glucose depletion, the mRNAs were strongly induced in the WT, but this was much less marked in *pin4Δ* or *pin4_ΔRRM_*(Fig. 6B). In the *pin4_RRM-M1_* strain induction of *HSP12* was reduced to a similar extent to *pin4Δ*, while induction of *HXT7* and *MTH1* was even lower in *pin4_RRM-M1_*.

The Pin4 C-terminal region is intrinsically disordered and potentially prion-forming (Calabretta and Richard 2015; Ottoz and Berchowitz 2020; Chowdhury et al. 2023), but includes the R3H domain. To test its function, we created a construct truncated after the N-terminal 200 aa (*pin4_ΔCT_*), fused with GFP to improve stability (Fig. 6A). Expression of Pin4_ΔCT_-GFP was similar to WT Pin4 (Supp. Fig. 9C). Following transfer of *pin4_ΔCT_* to Gly/EtOH for 16 min, transcriptional induction of *MTH1*, *HSP12*, and *HXT7* was significantly reduced relative to the WT, and very similar to *pin4Δ* and *pin4_ΔRRM_* (Fig. 6B). Statistical significance is shown in Fig. 6C.

### Pin4 is required for stress-induced accumulation of cytoplasmic granules

In parallel with the transcriptional response, glucose depletion in wildtype yeast induces formation of cytoplasmic granules over a minute time scale. Prominent granules contain Hsp104, an HSP100-family disaggregase that resolves stress-induced protein aggregates (Shorter and Southworth 2019). In live cell imaging of wildtype cells (Fig. 7A), 18% of cells had at least one Hsp104 granule on glucose medium, rising to almost 40% during the first 24 min following transfer to Gly/EtOH medium. Granule frequency plateaued around 70% after 90 min (Fig. 7B), as reported (Sathyanarayanan et al. 2020). In contrast, *pin4_RRMΔ_* showed a significantly reduced response. The percentage of cells exhibiting at least one Hsp104 granule increased only mildly, from approximately 20% in glucose to a plateau of around 30%, after 45 min in Gly/EtOH medium, and did not increase further (Fig. 7C). This muted response was also seen for total Hsp104-GFP induction (Fig. 7B, C).

**Figure 7:**
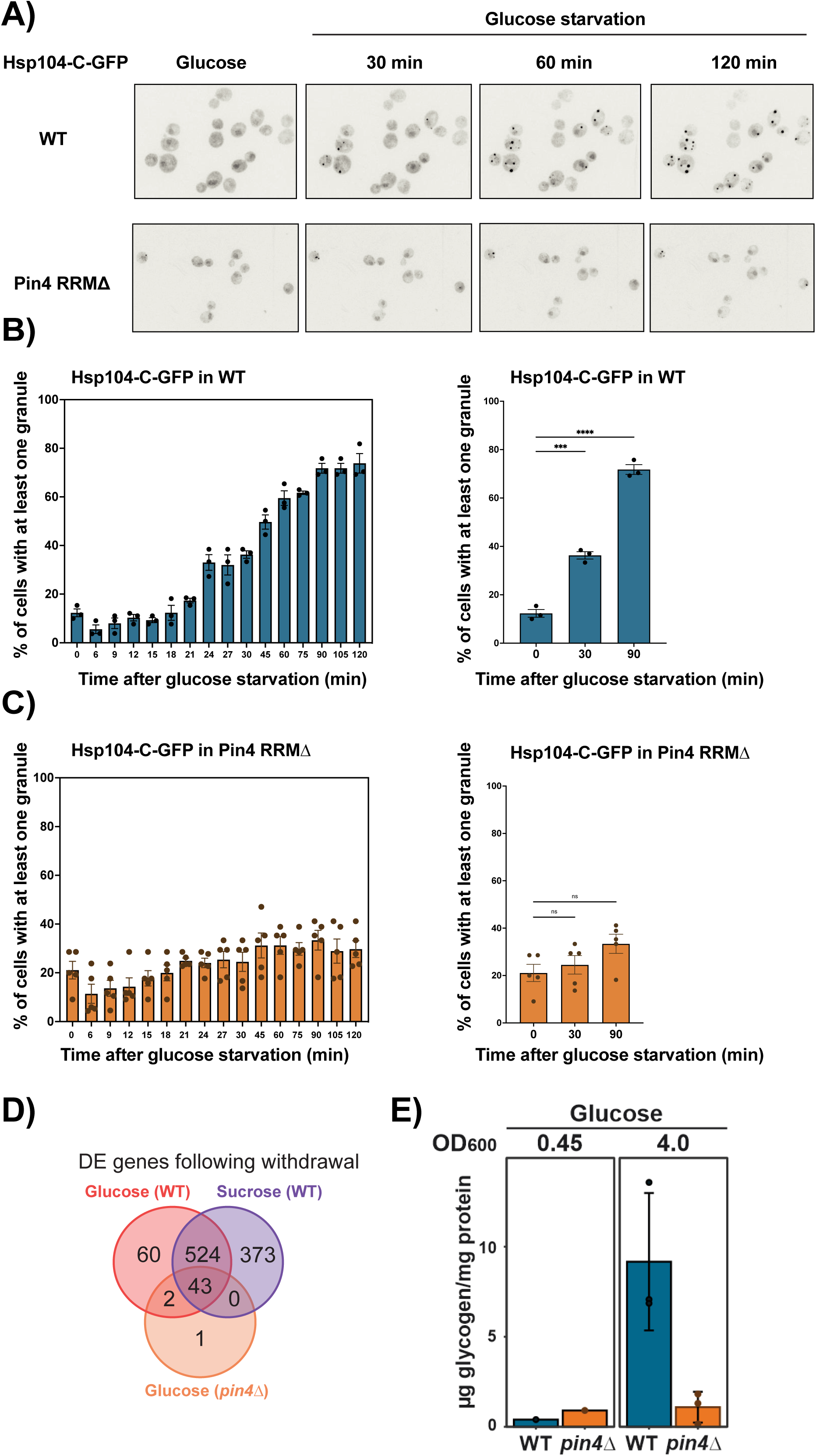
Loss of Pin4 or the RRM impairs energy stress responses A) Microscopy images showing the expression of Hsp104-GFP for WT and pin4*_RRM_Δ* strains in glucose and following glucose depletion for 30, 60, or 120 min. B, C) Bargraphs showing the percentage of cells with visible Hsp104-GFP foci in WT and pin4*_RRM_Δ* strains over a time course following glucose depletion. Blue bars represent the temporal dynamics of Hsp104 granule formation in WT cells, and orange bars in pin4*_RRM_Δ* cells. Statistical significance is indicated (*** p < 0.001, **** p < 0.0001, ns = not significant). Ordinary one-way ANOVA with Dunnett’s multiple comparisons test was used for the statistical test. Mean and individual replicate values are plotted. D) Venn diagram showing the overlap of transcripts significantly altered (fold-change > 2, false-discovery rate < 0.05) following withdrawal of either glucose or sucrose in wild-type (WT) cells, and glucose in *pin4Δ* cells. E) Quantification of glycogen levels in wild-type and *pin4Δ* yeast strains grown in SD-Trp to OD_600_ 0.45 (n= 1) or OD_600_ 4.0 (n=3).

We conclude that loss of either a functional RRM domain or the disordered C-terminal region of Pin4 blocks the transcriptional, and post-transcriptional, response to glucose withdrawal, suggesting functional interactions with both RNA and proteins.

### Pin4 acts between NTP-remodeling and transcriptional responses

We recently reported that glucose withdrawal is followed by a very rapid metabolic response, with NTP depletion over the initial 30 sec (Bexley et al. 2025). This results in the inhibition of translation initiation relative to elongation, leading to polysome collapse (Ashe et al. 2000; Bresson et al. 2020; Bexley et al. 2025). The relationship between NTP depletion and subsequent transcriptional adaptation is currently unclear. We considered that the loss of transcriptional adaptation in *pin4*Δ might lie downstream of an altered metabolic response.

However, glucose withdrawal from *pin4Δ* strains, induced rapid metabolic remodeling with depletion of intracellular ATP (Supp. Fig. 10A) and polysome collapse (Supp. Fig. 10B), similar to the wildtype.

We conclude that Pin4 is dispensable for the rapid, initial shutdown of protein synthesis following starvation.

### Pin4 is required for the response to energy depletion

Yeast, and many other organisms, have glucose-specific signaling pathways (reviewed in (Miles et al. 2021)). We therefore assessed whether the loss of transcriptional response in *pin4Δ* is specific for glucose removal, or a more general defect in responding to energy deficiency. Glucose-grown wildtype yeast cells predominately generate ATP through glycolysis (“aerobic fermentation”), whereas the yeast strains used here grown in sucrose medium generate ATP via hybrid mechanism of oxidative phosphorylation (“respiration”) and fermentation (Bexley et al. 2025). Common gene expression changes following withdrawal of either carbon source are likely responding to reduced energy availability, rather than a specific signaling pathway. Wildtype cells growing on 2% sucrose medium, were energy limited through rapid transfer to 2% Glyc/EthOH. 567 genes showed altered expression following both glucose and sucrose withdrawal. Only 43 (8%) of these were altered in the *pin4Δ* strain following glucose withdrawal (Fig. 7D). We conclude that transcriptional changes linked to general energy depletion are suppressed in the absence of Pin4.

In wildtype yeast, glycogen storage is reported to be triggered by decreased energy availability as cells leave log phase growth and approach quiescence (reviewed in (Miles et al. 2021)). In glucose medium, glycogen storage begins when about half the glucose has been taken up from the medium. We therefore measured glycogen accumulation in WT and *pin4Δ* cells, at mid-log (OD_600_ 0.45) and a later growth stage (OD_600_ 4.0). In the mid-log samples, corresponding to the growth stage used for nutrient shift experiments, glycogen was scarcely detected above background in either the WT or *pin4Δ* cells. In the samples at higher OD, glycogen strongly accumulated in the wildtype, but this was greatly reduced in *pin4Δ* cells (Fig. 7E). We conclude that Pin4 is also required for the response to reduced energy availability during normal growth.

We conclude that following energy depletion Pin4 acts between initial NTP-remodeling and subsequent transcriptional, and post-transcriptional responses (Fig. 8).

**Figure 8:**
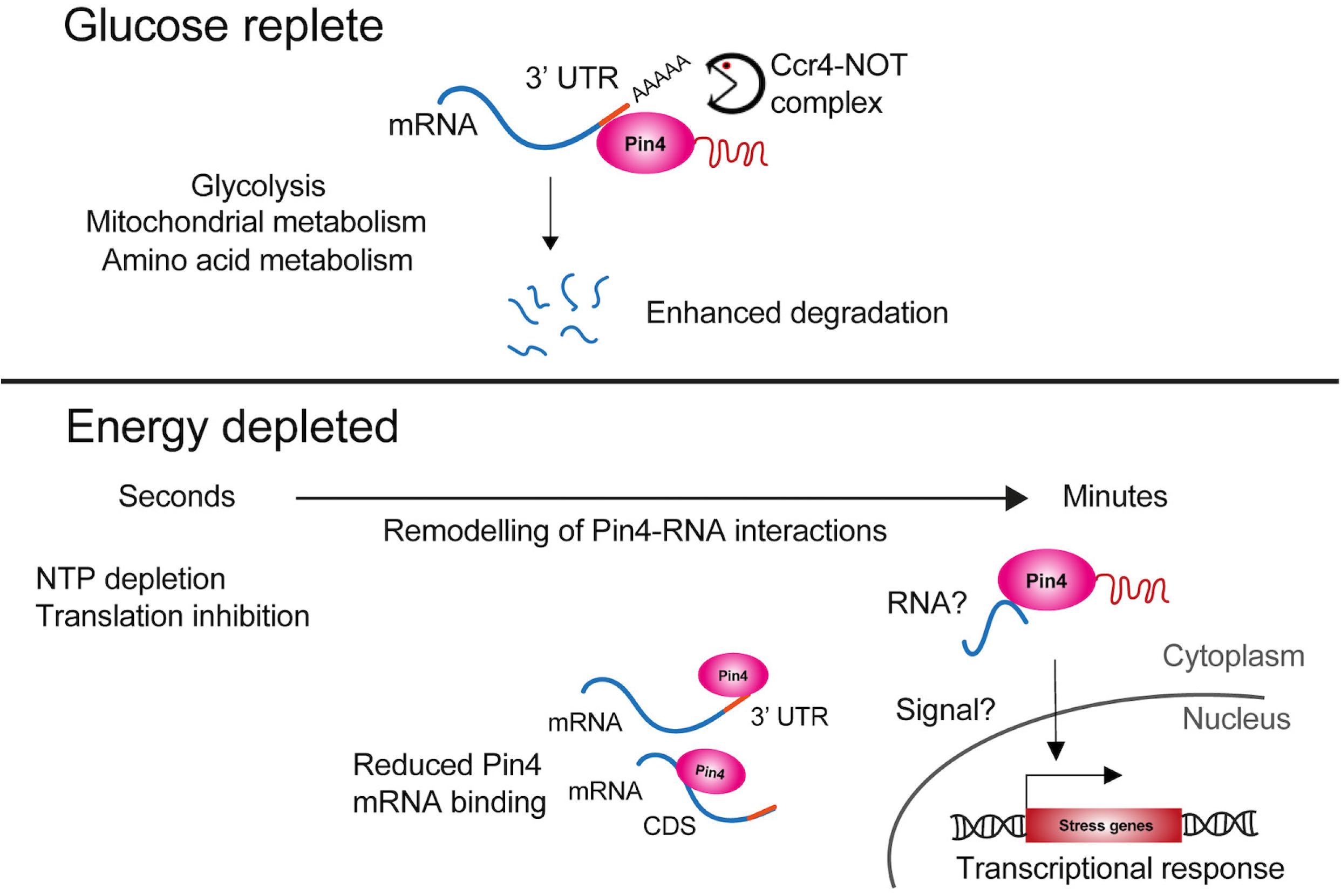
Model for Pin4 functions Upper panel: In glucose replete conditions, Pin4 binds to the 3’ untranslated regions (UTRs) of mRNAs. Preferentially those encoding glycolytic enzymes, and components of mitochondrial and amino acid metabolism. This promotes mRNA destabilization, potentially via interactions with components of the Ccr4-NOT complex. Lower panel: Upon energy depletion NTPs deplete within seconds, independently of Pin4. Interactions between Pin4 and RNA are remodeled, greatly reduced mRNA binding. Pin4 is also needed to signal the transcriptional reprogramming, although the precise role of Pin4 in signaling remains unclear. The intrinsically disordered region is needed for this response, as is a functional RRM. We propose that the transcriptional stress response is subject to riboregulation, but the specific RNA required remains unclear.

## DISCUSSION

RBPs play crucial roles in cellular responses to many environmental stresses. We previously screened for global changes in the RNA-protein interactome on a short timescale following a variety of stresses, including heat shock and glucose withdrawal (Bresson et al. 2020).

Unexpectedly, a set of largely uncharacterized proteins were among the most stress responsive RBPs. Included in these “unknome” factors was Pin4, which we functionally characterized here.

Using the newly developed reCRAC technique (Ristová et al. 2025), Pin4 showed a distinctive pattern of binding to mRNA 3’ UTRs during growth on glucose medium. The mRNAs with strong Pin4 binding, relative to RNA abundance, all carried a clear consensus motif. This was located close to the site of crosslinking, which can be identified at the nucleotide level in reCRAC, strongly supporting its role as a direct Pin4 recognition element. However, 3’ UTRs of almost all mRNA were detectably bound by Pin4, indicating that recognition by Pin4 has at least two components: Recognition of some feature(s) common to all mRNA 3’ UTRs, plus binding to a specific sequence motif. Notably, the functional classes of mRNAs favored for Pin4 binding were not random. They included most glycolytic enzymes, plus several key nuclear-encoded mitochondrial proteins and a small number of important enzymes in amino acid metabolism.

At short time points following glucose depletion, mRNA levels were not substantially altered as previously reported (Zid and O’Shea 2014; Bresson et al. 2020), but overall crosslinking of Pin4 to mRNAs was substantially reduced. However, the preference for 3’ UTR binding and the same enriched motif were preserved, suggesting that the recruitment mechanism is retained. The reduced interactions could reflect altered Pin4 affinity, e.g. due to phosphorylation, which is altered at multiple sites in response to stress (Smolka et al. 2007; Soufi et al. 2009; Gnad et al. 2009; Soulard et al. 2010; Bodenmiller et al. 2010; Helbig et al. 2010). Alternatively, Pin4 may form aggregates via its prion-like domain (Derkatch et al. 2001; Yang et al. 2013, 2014) and/or mRNA availability might be reduced by sequestration into cytoplasmic condensates (Teixeira et al. 2005; Glauninger et al. 2024). On translation-related transcripts, residual Pin4-binding was seen across the CDS. Following glucose withdrawal, these pre-stress mRNAs are still present in large quantities but are largely ribosome-free (Zedan et al. 2024). The CDS signal might reflect colocalization with Pin4 in the absence of sequence-specific binding, potentially within cytoplasmic condensates (Teixeira et al. 2005; Glauninger et al. 2024).

Analyses using reporter constructs indicated that Pin4 binding modestly destabilized most, but not all, interacting mRNAs, under both glucose-replete and glucose-depleted conditions, with consequent reductions in reporter protein accumulation. This conclusion was supported by RNA sequencing in strains lacking Pin4, and a recent, systematic analysis of protein abundances in yeast deletion mutants (Messner et al. 2023). Previous high-throughput analyses reported protein-protein interactions between Pin4 and components of the CCR4- NOT complex (Caf40, Cdc39, Cdc36, Mot2, Ccr4), which mediates mRNA degradation (Tarassov et al. 2008; Schlecht et al. 2012; Miller et al. 2017). Notably, the abundances of Ccr4 and other components of the CCR4-NOT complex, are very similar to that of Pin4. We postulate that transient Pin4 binding within the 3’ UTR can promote mRNA degradation through increased association of the CCR4-NOT complex. Retargeting of Pin4 onto non-ribosome bound, translation-related transcripts following stress would fit this model.

The preferred mRNA targets for Pin4 in glycolysis and mitochondria suggested roles in energy metabolism. In glucose medium, *pin4Δ* cells exhibited hypersensitivity to both H_2_O_2_, which induces oxidative stress, and glyoxal, a reactive carbonyl species that generates reactive oxygen species and disrupts mitochondrial membrane potential (Shangari and O’Brien 2004). We speculate that this hypersensitivity is linked to misregulated mitochondrial function due to imbalances in protein production in the absence of Pin4.

Unexpectedly, loss of Pin4 profoundly affected transcriptional adaptation to glucose depletion, almost completely abolishing the induction of stress-responsive mRNAs. We identified 645 mRNAs that were significantly differentially expressed between Pin4-deficient cells and WT cells following starvation. To test the requirement for RNA binding we generated strains lacking the single RRM of Pin4 or carrying RRM point mutations predicted to disrupt protein-RNA binding but not domain structure. Both mutants recapitulated the *pin4Δ* transcription phenotype in response to glucose withdrawal. Deletion of C-terminal region in *pin4_ΔCT_* also blocked transcriptional reprograming. The region deleted was previously reported to form prion-like aggregates (Derkatch et al. 2001). However, initial live cell imaging did not reveal clear Pin4 aggregation on glucose withdrawal.

Comparison to wildtype subjected to sucrose withdrawal, indicated that *pin4Δ* cells fail to respond to energy deficiency, rather than specific glucose depletion. Consistent with this model, *pin4Δ* cells also failed to induce glycogen storage when approaching stationary phase. In the wildtype,Hsp104 granules form in response to protein misfolding following environmental insults (Uvdal and Shashkova 2023). Live cell imaging revealed that *pin4_ΔRRM_* cells almost entirely failed to induce Hsp104 granule formation, highlighting the loss of stress response. Given their lack of response to energy depletion, it was unsurprising that *pin4Δ* cells displayed greatly impaired recovery of growth following transfer from glucose to non-fermentable carbon sources.

In contrast, cells lacking Pin4 retained the initial (second time scale) metabolic response to glucose withdrawal, during which ATP is very rapidly depleted leading to loss of translation initiation. Metabolomic remodeling, which drives the initial translational response to energy depletion, is therefore independent of Pin4.

In summary, we propose a dual-function model for Pin4 in glucose-rich and glucose-depleted conditions (Fig. 8). In glucose, Pin4 binds to the 3’ UTRs of metabolic and pro-growth mRNAs, and regulates gene expression post-transcriptionally by promoting their destabilization. Following energy depletion, Pin4 is required to activate the transcriptional stress response through mechanisms involving its RNA recognition motif and prion-forming domain. Previous genetic screens for glucose signaling pathways did not identify Pin4, but these were based on timescales of days. In contrast, a timescale of minutes is optimal for rapid and reversible responses to environmental stimuli. Loss and restoration of nutrient availability to yeast growing on plant surfaces might occur several times a day, and transient but substantial stresses will be the norm in many biological systems. Post-transcriptional mechanisms likely provide the first responses, on timescales of seconds to minutes, followed by longer term transcriptional changes that can persist for days. We propose a bridging role for Pin4, connecting an initial metabolic shift to subsequent transcriptional responses (Fig. 8). The requirement for RRM function further suggests that this activity is riboregulated.

## LIMITATIONS OF THE STUDY

The exact point at which Pin4 operates within the glucose signaling pathways, and how its absence blocks the transcriptional response, remains unclear (Fig. 8). Previous data suggest links between glycolytic flux and signaling (Conrad et al. 2014), but the mechanistic link is not defined and Pin4 might function here.

Our *pin4Δ* strains are steady state, as opposed to sudden loss of Pin4, and some changes observed could be due to secondary effects. It might be informative to test the effects of acute Pin4 depletion, to better assess whether the presence of Pin4 prior to, or immediately following, glucose deprivation is required for the transcriptional response.

The analyses indicate that RNA interactions may be important for the transcriptional role of Pin4, but the nature of the RNA species involved remains unclear (Fig. 8).

## MATERIALS AND METHODS

### Reagents

10 mM dNTP mix (Thermo Scientific, Cat#18427013) 10x NEBuffer 2.1 (NEB, Cat#B7202)

10x T4 DNA ligase buffer (NEB, Cat# B0202S)

5-alpha Competent E.coli High Efficiency (NEB, Cat# C2987I) Act1 primary antibody (Abcam, Cat# 8224)

BamHI-HF (NEB, Cat# R3136S) BclI (NEB, Cat#R0160S)

BsaI-HF-v2 (NEB, Cat# R3733S) BsmBI-v2 (NEB, Cat# R0739S)

Concanavalin A (Sigma, Cat# C2272-2MG) CSM Complete (Formedium, Cat#DCS0011)

CSM Single Drop-out -Trp (Formedium, Cat#DCS0149)

Cycloheximide – stock of 20 mg/ ml in EtOH prepared and stored at -20C (Sigma-Aldrich, Cat#C7698-5G)

D (+) – Glucose Anhydrous (Thermo Scientific, Cat#10141520) DNase/RNase free water

EcoRI-HF (NEB, Cat# R3101S)

EDTA-free cOmplete protease inhibitor cocktail (Roche, Cat#11836170001) FLAG antibody (Sigma Aldrich, Cat# SAB4200071)

Glyoxal solution (Merck, Cat#128465) HindIII-HF (NEB, Cat# R3104S)

Hydrogen Peroxide (Acros Organics, Cat#7722-84-1) Klenow exo- (NEB, Cat#M0212L)

Microscopy chamber with 8 individual wells high glass bottom (Thistle Scientific, Cat# IB- 80807)

MOPS running buffer 20x (Invitrogen, NP0001) Nitrocellulose membrane (Thermo Scientific, Cat#88018) NotI-HF (NEB, Cat# R3189S)

NuPAGE Bis-Tris 4-12% precast gradient gels (Invitrogen, Cat#NP0321BOX) OneShot Top10 Chemically competent E. coli (Thermo Scientific, Cat#C404003) Phusion High-Fidelity DNA polymerase (NEB, Cat# M0530S)

Random hexamers (Thermo Scientific, Cat# 10609275) RNasin (Promega, Cat# N2511)

RQ1 DNase (Promega, Cat# M6101)

Superscript III RT, 0.1M DTT, 5x First Strand Buffer (Invitrogen, Cat#18080093) SwaI (NEB, Cat#R0604S)

T4 DNA ligase (NEB, Cat# M0202L)

TRIzol Reagent (Invitrogen, Cat#15596026)

Venus primary antibody (Agrisera, Cat# AS184227) Yeast Nitrogen Base (Formedium, Cat#CYN0410) Zirconia beads (Thiste Scientific, Cat#ZrOB05)

### Commercial kits

Gel extraction kit (QIAGEN, Cat#28704)

Min Elute PCR purification kit (QIAGEN, Cat#28004) Plasmid miniprep (ZYMO Research, Cat# D4016)

RNA clean & concentrator kit (ZYMO Research, Cat# R1013) Zymoclean Gel DNA recovery kit (ZYMO Research, Cat# D4007)

**Software and Algorithms** BEdtools (Quinlan and Hall 2010) EdgeR (Robinson et al. 2010) Flexbar (Dodt et al. 2012) ggplot2 (Wickham 2016)

IGV (Robinson et al. 2011)

Metamorph software (Molecular Devices) Novoalign (V2.07.00, Novocraft Technologies) GraphPad Prism Version 10.0.3 Proteomaps(Liebermeister et al. 2014) pyCRAC (Webb et al. 2014)

PyMOL 3.1(Molecular Graphics System) Salmon(Patro et al. 2017)

### Data Availability

#### Oligonucleotides

All oligonucleotides used in this study are listed in Supplementary Table 1.

#### Strains

All strains generated in this study are listed in Supplementary Table 2 and are available from the lead contact upon request.

#### Plasmids

All plasmids generated in this study are listed in Supplementary Table 3 and are available from the lead contact upon request.

#### Deposited data

All reCRAC and RNAseq data are included in the GEO accession: https://www.ncbi.nlm.nih.gov/geo/query/acc.cgi?acc=GSE283345.

## METHOD DETAILS

### Yeast cell culture and media

All Saccharomyces cerevisiae strains used in this paper originated from the BY4741 background (MATa his3Δ1 leu2Δ0 met15Δ0 ura3Δ0). Yeast cultures were grown in a growth medium containing 2% D-glucose, 2% yeast nitrogen base, and a 2% complement of amino acids excluding tryptophan (SD-Trp) at 30°C shaking. For all the experiments, the cultures were grown to a stationary phase overnight and then diluted the next morning from OD_600_ of 0.05 to OD_600_ of 0.45. For experimental treatments, yeast cells were either harvested directly from glucose-supplemented media (Glucose) or subjected to glucose deprivation by transferring to the same media but lacking glucose and instead supplemented with 2% glycerol and 2% ethanol (no Glucose). For the polysome profiling experiment, cells were grown in synthetic complete (SC) medium.

For reCRAC, cells were crosslinked with UVC at 254 nm at an OD_600_ of 0.45. Harvesting for both reCRAC, RNAseq and qPCR experiments involved rapid filtration of the cultures onto a nitrocellulose membrane, immediate immersion of the membrane into 50 mL of ice-cold water, centrifugation at 4,600 rpm for 2 min at 4°C, and subsequent freezing of the cell pellet.

### Gene tagging, mutations and deletion

For reCRAC experiment, the chromosomal locus of Pin4 was modified to include an N-terminal FLAG-His (FH) tag. This tag consists of a FLAG sequence (GATTATAAAGATGACGATGACAAG), a four-alanine spacer (GCCGCAGCCGCA), and a chain of eight histidine residues (CATCACCACCATCATCACCATCAC).

The integration of the FH tag was done using a CRISPR-Cas9 system, utilizing the pML104 plasmid as described (Laughery et al. 2015). This plasmid is engineered to include the Cas9 sequence, an ampicillin resistance gene, and the URA3 selection marker. It is also optimized for efficient and directional cloning of single guide (sg) RNA. The design features an sgRNA with a GATC overhang compatible for cloning, which is a 20-nucleotide guide sequence that targets genomic DNA (e.g. the N-terminus of Pin4), and a structural segment critical for Cas9 binding (see Supp. Table 1). The yeast CRISPR-Cas9 system stimulates homologous recombination using a repair template that is co-transformed with the Cas9 and sgRNA expressing vector. This template includes homology regions close to the cleavage site, the sequence to be introduced/changed, and mutations either in the sgRNA target or in the adjacent PAM sequence (NGG) to confer resistance to further Cas9 cleavage in modified cells (see Supp. Table 1).

Briefly, for sgRNA preparation,10 µg of pML104 plasmid was digested overnight at room temperature with 2 µL of SwaI, subsequently inactivated at 65°C for 20 min, and further digested with 2 µL of BclI at 50°C for 2 hours. Following gel extraction with QIAGEN Gel extraction kit, the cut plasmid was ethanol precipitated, resuspended in water, aliquoted and stored at -80°C. The sgRNA oligos (Supp. Table 1) were annealed by combining 1 µM of each oligo with 1x T4 DNA ligase buffer in a total volume of 100 µL, heated at 95°C for 5 min, and then allowed to cool gradually to 20°C with the speed of 45s/°C. The annealed oligos were ligated with 66 ng of linearized pML104 plasmid using 0.5 µL T4 DNA ligase and 1x T4 DNA ligase buffer in a reaction volume of 10 µL. The mixture was kept at room temperature for 2 hours, transformed into OneShot Top10 Chemically Competent E. coli cells, and plated on LB-Amp plates. Plasmid DNA was isolated and sequenced to confirm the accurate sgRNA insertion in pML104 plasmid.

The FH tag was integrated using two repair templates consisting of the FH sequence flanked by 45 bp homology arms with engineered silent mutation to disrupt the PAM site, and 20 bp overlap at their 3’ ends (Supp. Table 1). The repair template was made by annealing two 10 µM repair template oligos in 1x NEBuffer 2.1 in a reaction volume of 50 µL at 95°C for 6 min, and then decreasing the temperature by 1°C every 30s until 20°C was reached. The repair template was then extended with 1 µL of Klenow exo- together with 1.25 µL of 10 mM dNTPs for 1 hour at 37°C. 10 µL of the annealed and extended repair template was then co-transformed with 500 ng of sgRNA containing-pML104 plasmid into BY4741 strain using the standard PEG/LiOAc yeast protocol. Selection occurred twice on -URA plates, with further restreaking on YPD to remove the selection plasmids. Colonies were then restreaked on both YPD and -URA, and the ones that lost the pML104 plasmid and failed to grow on -URA were verified by PCR using the check oligos (Supp. Table 1) and sequenced for the correct insertion of the FH tag.

The pin4*_RRMΔ_* strain, which involves deletion of the entire RRM domain from A86 to V158, and the pin4_RRM-M1_ strain (V88A, L128A, F130A) was generated in the N-FH Pin4 background using the CRIRPS/Cas9 system, following the protocol described above. See Supp. Table 1 for the used oligos. For the complete deletion of the Pin4 gene, we employed conventional homologous recombination, utilizing the KanMX6 cassette from the pFa6a plasmid (Addgene #39296), with oligonucleotides detailed in Supp. Table 1.

The Hsp104 protein was C-terminally tagged with GFP (S65T version) using a PCR-based method from the pFa6A plasmid (Addgene #39293), incorporating the KanMX6 selection marker (see Supp. Table 1). The resulting construct was transformed into the BY4741 strain and pin4*_RRMΔ_* strain. The pin4*_ΔCT_*, where GFP was inserted after the first 200 amino acids of Pin4 was made by taking the GFP (S65T) from the pFa6A plasmid (Addgene #39293) and using it as the repair template utilizing CRISPR-Cas9 system as described above.

### Growth curves

For the growth curves, yeast strains were inoculated from single colonies and grown overnight in SD-Trp medium. The cultures were then diluted to an optical density OD_600_ of 0.05 and grown to an OD_600_ of 0.45 in SD-Trp. At this exponential growth phase, cultures were diluted to an OD_600_ of 0.025 and 120 µL aliquots were pipetted into a 96-well plate. Stress-inducing agents were added to the wells at varying final concentrations: glyoxal at 0.3, 1,5, 3, 9 and 30 mM; and hydrogen peroxide at 0.5 and 1 mM. For Antimycin A sensitivity assays, cultures were saturated overnight in SD medium. Subsequently, they were diluted to OD600 0.05 in 20 ml of fresh media and grown to an OD_600_ of 0.3. The cells were then shifted to SD medium without any supplement or supplemented with 0.2 or 0.02 µg/ml of Antimycin A. The 96-well plate was sealed with a sealing tape to minimize evaporation and placed in a Sunrise TECAN plate reader with a continuous shaking at 30°C. Optical density measurements were taken every 15 min to monitor growth. Each condition was tested in technical triplicate and biological duplicate. Specific strains are provided in the corresponding Fig. legends.

For the glucose starvation experiment, overnight saturated BY4741 and *pin4Δ* cells were diluted to an OD_600_ of 0.05 and grown to an OD_600 of_ 0.40 in SD-Trp. Cells were then centrifuged, washed, resuspended in a medium containing 2% glycerol and 2% ethanol, and grown in this medium from an OD_600_ of 0.075. Growth was monitored over a period of 2.5 days. This condition was tested in a biological duplicate.

### reCRAC

See Ristová et al,. (2025) for a detailed protocol and Fig. 1B for a visual overview of reCRAC. Briefly, Pin4 was engineered to include an N-terminal FLAG-4xAlanines- 8xHistidines (FH) tag expressed from its native promoter to ensure it is the only variant produced. Expression of the tagged construct was verified by western blot (Supp. Fig. 3A). The strain was grown in SD-Trp, and upon reaching the exponential growth, cultures were UV crosslinked at 254 nm to stabilize RNA-protein interactions either in glucose medium or shifted to glucose-deprived medium (2%glycerol, 2% ethanol) for 16 min. xposure to UV irradiation was performed for only 8 sec and all covalent RNA-protein interactions are generated during this period. We do not expect a significant stress response to develop during this very short interval. Following irradiation cells are immediately cooled and harvested. Changes in RNA-protein interactions that occur during harvesting and lysis will not affect recovery of RNA-protein interactions in reCRAC.

The lysis step involved highly denaturing conditions with 6 M Guanidine – HCl to prevent protein degradation. These lysis conditions are very similar to the Trizol lysis used for RNAseq analyses. Subsequent purification and wash steps were performed on Ni-NTA beads under similarly stringent conditions. The adapter ligation, radiolabeling with ^32^P and partial Benzonase digestion to generate protein-protected RNA fragments were performed on the beads. Following elution from the Ni-NTA beads, a second purification on anti-FLAG beads was performed under more native conditions. The samples were then visualized via autoradiography, the RNA bound to the target protein was cut out and treated with Proteinase K. The RNA was then converted to cDNA, amplified and analyzed by high-throughput sequencing.

### Reporter constructs

To study the effect of Pin4 binding on its RNA targets, we made reporter constructs using the modular cloning system (MoClo) (Lee et al. 2015). This system involves assembling approximately nine modular parts (Part plasmids) – including different promoters, coding regions, terminators, etc. – in a manner similar to LEGO, creating a complete Cassette reporter plasmid in a single golden gate assembly reaction, which can be chromosomally integrated.

All toolkit Part plasmids except the terminator Part plasmids used in this study were obtained from Addgene and are found in Supp. Table 3. The terminator Part plasmids containing the sequences of 3’ UTRs of Pin4 reCRAC targets (TPI1, GPM1) and a non-target control (PNC1) were generated for this study using PCR and standard cloning techniques.

The process to generate the terminator Part plasmids began by modifying the target terminator sequence to incorporate BsmBI and BsaI restriction sites via PCR amplification. This was done by extracting DNA from the parental By4741 strain using 100 µL of 200 mM LiOAc and 1% SDS solution. The sample was incubated at 70°C for 5 min, then precipitated with ethanol. The purified DNA then served as the template for PCR reactions using Phusion DNA Polymerase, with oligonucleotides listed in Supp. Table 1 (oligos #526, 527, 528, 529, 532, 533). We verified the PCR products on a 1% agarose gel, excised the appropriate bands, and purified the fragments using the Zymoclean Gel DNA Recovery Kit.

200 ng of the purified PCR products were digested with 0.5 µL of BsmBI-v2 in a total reaction volume of 40 µL for 2h at 55°C. The restriction enzyme was then heat deactivated at 80°C for 20 min. The reaction products were subsequently purified using the Min Elute PCR Purification Kit. We then ligated 80 ng of the digested inserts with 65 ng of the similarly treated empty entry vector (pYTK001) using 0.5 µL of T4 DNA ligase for 1.5h at room temperature. The ligase was inactivated by putting the samples at 65°C for 10 min. The terminator Part plasmids were then transformed into NEB 5-alpha E. coli competent cells and plated onto LB- 25 µg/ml Chloroamphenicol plates, purified and verified by sequencing.

For assembling the complete Cassette plasmid, we combined individual Part plasmids in a reaction containing 74 ng of each part plasmid, 1 µL T4 DNA ligase, 2 µL T4 DNA ligase buffer, 1 µL BsaI in 20 µL volume reaction (see Supp. Table 3 for the detail of specific Part plasmids). The samples were incubated in a thermocycler with the setting of 25 cycles of 42°C for 2 min, 16°C for 5 min, with a final digestion step (60°C for 10 min) and heat inactivation (80°C for 10 min). The PCR products were then transformed into NEB 5 alpha E. coli competent cells. The assembly was verified by diagnostic digest using EcoRI-HF, NotI-HF, BamHI-HF, HindIII-HF restriction enzymes. Finally, the confirmed Cassette plasmids were linearized with NotI for 10 min at 37°C, transformed into target yeast strains (By4741, *pin4Δ*), which we plated on YPD plates with 600 µg/ml hygromycin. The integration into the chromosome was confirmed by PCR and sequencing using oligos listed in Supp. Table 1.

### Western blot analysis of reporter constructs

For the analysis of the Venus reporter expression via western blot, cells expressing Venus reporters were initially grown overnight in SD-Trp medium. The following morning, the cultures were diluted to an OD_600_ of 0.05 and grown until reaching an OD_600_ of 0.45. Cells corresponding to 9 ODs were harvested by centrifugation at 4,600 rpm for 2 min and subsequently frozen at -90°C for storage. For glucose starvation experiments, cells were rapidly filtered and resuspended in a medium containing 2% glycerol and 2% ethanol, where they were incubated for 2 hours before spinning them and freezing them the same way as the non-starved samples.

Cell pellets were resuspended in 1 mL of 0.2 M NaOH and incubated on ice for 10 min. Subsequently, 100 µL of 50% trichloroacetic acid (TCA) was added to precipitate proteins. After mixing, samples were incubated on ice for an additional 10 min and then centrifuged at 14,000 rpm at 4°C for 10 min to pellet the precipitated proteins. The pellet was resuspended in 35 µL of lysis buffer (0.1 M Tris-HCl pH 6.8, 4 mM EDTA pH 8.0, 4% SDS, 20% glycerol, 2% β-mercaptoethanol). To adjust the pH, 15 µL of 1M Tris base was added to each sample. Samples were then heated at 95°C for 10 min and centrifuged at 10,000 rpm at room temperature for 10 min. The supernatant was kept, and the protein concentration was quantified using the Bradford assay. 25 µg of protein per sample was loaded onto a 4-12% Bis-Tris gel and subjected to electrophoresis at 180V for 1 hour in MOPS buffer. Proteins were then transferred to a nitrocellulose membrane using the i-blot2 system set on the P0 setting. Post-transfer, the membrane was stained with Ponceau S solution for 5 min to visualize total protein transfer and subsequently blocked with 5% milk in PBS for 30 min. The membrane was incubated with primary antibodies against Venus (Agrisera AS18 4227, diluted in 1:1000) and Act1 (Abcam 8224, diluted 1:10,000) for 1 hour. After washing three times for 5 min each with PBS-T (0.1% tween), the membrane was incubated with a fluorescently labelled secondary antibody (LiCor, diluted 1:5,000 in 5% milk PBS-T) for 1 hour. Following three additional washes with PBS-T (0.1% tween), the membrane was scanned using the LI-COR software (LI-COR Biosciences). Act1 served as the loading control, the quantification was done using the ImageStudio (LI-COR Biosciences).

### RNAseq

BY4741 and *pin4Δ* cells were cultured in 100 mL of SD-Trp to OD600 of 0.45 from OD600 of 0.05. Cells were rapidly filtered and immediately immersed in 50 mL of ice-cold water, centrifuged at 4,600 rpm for 2 min at 4°C, and the cell pellet was snap-frozen in a dry-ice and ethanol bath, and stored at -80°C. For glucose starvation stress, cells were incubated in SD-Trp medium without glucose but supplemented with 2% glycerol and 2% ethanol. At the 16 min timepoint, cells were filtered and immersed in 50 mL of ice-cold water, after which they were spun down as described for glucose samples, and snap-frozen. Biological triplicates were collected for each condition. For sucrose experiment, BY4741 cells were grown to OD600 0.45 in 50 ml SC-Trp media supplemented with 2% sucrose.

RNA was extracted from 10 OD_600_ units of cells using TRIzol reagent with the addition of 0.5% final β-mercaptoethanol. Cells were lysed by bead beating in screw-cap tubes three times in a Fast Prep-24 machine (MP Biomedicals) using 200 µL of Zirconia beads at a setting of 6 m/s for 40 sec. RNA was cleaned twice by chloroform extraction and precipitation was performed using 0.8 vol isopropanol. RNA concentration was quantified using Nanodrop (OD_280_). 6 µg of RNA was treated with DNase I for 30 min and further purified using RNA Clean and Concentrator kit.

Libraries for RNAseq were prepared by the Edinburgh Clinical Research Facility at the Institute of Genetics and Cancer (Edinburgh, UK). RNA integrity and quality were assessed using an Agilent 2100 Electrophoresis Bioanalyser Instrument (Agilent Technologies Inc, #G2939AA) with RNA 6000 Nano chips (#5067-1511). RNA quantification was performed with Qubit 2.0 Fluorometer (Thermo Fisher Scientific Inc, #Q32866) using the Qubit RNA broad range assay kit (#Q10210). DNA contamination levels were evaluated using the Qubit dsDNA HS assay kit (#Q32854).

Libraries were prepared from 500 ng of total RNA per sample using the NEBNEXT Ultra II Directional RNA Library Prep kit (NEB #7760) and the Poly-A mRNA Magnetic Isolation Module (NEB #E7490) following the manufacturer’s protocols. Sequencing was carried out on the NextSeq 2000 platform (Illumina Inc, #20038897) using NextSeq 2000 P3 Reagents (200 Cycles) (#20040560) with paired-end, 2x 100 nt outputs.

### Quantitative PCR (qPCR)

Yeast strains BY4741, *pin4Δ*, pin4*_RRMΔ_*, pin4*_RRM-M1_*, and Pin4*_ΔCT_* were cultured overnight in SD-Trp medium. The next day, cultures were diluted to an OD_600_ of 0.05 and grown in 100 mL of SD-Trp to an OD_600_ of 0.45. For glucose starvation experiments, cells were filtered and resuspended in 100 mL of medium containing 2% glycerol and 2% ethanol for 16, 32, 60 or 480 min. For harvesting, cultures were rapidly filtered, and the cells were immediately immersed in 50 mL of ice-cold water, centrifuged at 4,600 rpm for 2 min at 4°C, and cell pellets were frozen.

For RNA extraction, cell pellets corresponding to 10 OD_600_ units were lysed using 100 µL of TRIzol reagent in the presence of 200 µL zirconia beads and 0.5% β-mercaptoethanol. The lysis was performed in screw-cap tubes twice using a fast prep machine at 6 m/s for 40 seconds. Extra 500 µL of TRIzol reagent with 0.5% β-mercaptoethanol were added and the bead beating was repeated one more time. The samples were cleared by centrifugation, and RNA was extracted using a chloroform-isopropanol method. The RNA was washed with 70-75% ethanol, dried and resuspended in DNase/RNase-free water. The RNA concentration was quantified using a Nanodrop spectrophotometer.

For reverse transcription, the extracted RNA was first treated to remove any contaminating genomic DNA. 15 µg of RNA was mixed with 7 µL of RQ1 DNase and 1x buffer, along with 1 µL of RNasin. This mixture was gently vortexed and incubated at 37°C for 30 min. The DNase was then inactivated by adding 125 µL of water to the mixture, followed by an extraction with equal volume of phenol:chloroform:isoamyl alcohol (25:24:1), vortexed vigorously, and centrifuged at 12,000g for 10 min at 4°C. The aqueous phase was extracted, treated with chloroform and isoamyl alcohol (24:1), and the RNA was precipitated overnight at -20°C using sodium acetate and ethanol. The RNA pellet was washed, dried, and dissolved in DNase/RNase-free water. The concentration and purity of RNA were assessed using a Nanodrop spectrophotometer.

Reverse transcription was performed on 1 µg of DNase-treated RNA using random hexamers and Superscript III Reverse Transcriptase (Invitrogen). The reaction mixture included 0.2 µg/µL random hexamers, 10 mM dNTP mix, 5x first-strand buffer, 0.1M DTT, RNasin, and reverse transcriptase. The mixture was incubated at 65°C for 5 min, followed by synthesis conditions of 25°C for 15 min, 55°C for 1 hour, and a final inactivation step at 70°C for 15 min. Finally, 180 µL of water was added to the reaction to obtain a total volume of 200 µL of cDNA.

Quantitative PCR was carried out in triplicate using SYBR Green I master mix (Roche) on an MX3005P qPCR system (Agilent Technologies). Each 10 µL reaction contained 4 µL of cDNA, 5 µL of 2x SYBR Green master mix, 0.4 µL of 10 µM primers (forward and reverse mixed at 1:1), 0.04 µL of Rox II and 0.56 µL water. The thermal cycling conditions were as follows: initial denaturation at 95°C for 30 seconds, followed by 38 cycles of 95°C for 5 seconds, 55°C for 15 seconds, and 72°C for 15 seconds.

### Glycogen Assay

Yeast strains BY4741 and *pin4Δ* were cultured overnight in SD-Trp medium to obtain saturated cultures. These cultures were then diluted in 100 mL of SD-Trp and grown to an OD_600_ of either 0.45 or 4.0. Cells were harvested by centrifugation, washed with 10 ml ice-cold water and transferred to screw-cap tubes. Cell pellets were resuspended in 300 µl ice-cold water and 200 µl zirconia beads were added. The cells were lysed by three rounds of beat beating with a FastPrep-24 (MP Biomedicals) machine set to 6 m/s for 40 sec with 1 min breaks on ice. Beads and cell debris were pelleted by centrifugation and the supernatant transferred to a new tube. The lysate was cleared by subsequent centrifugation steps at 13,000 rpm for 10 and 5 min at 4°C. The protein concentration was determined with the Bio-Rad Protein Assay Dye Reagent Concentrate (Bio-Rad, #5000006). The remaining lysate was boiled for 20 min to inactivate all enzymes. The denatured proteins were removed by centrifugation at 18000 xg for 10 min. The glycogen contained in the supernatant was measured with the Glycogen Assay Kit II (Colorimetric) (abcam, ab169558) in a SpectraMax M5 plate reader (MOLECULAR Devices). Each sample contained the volume corresponding to 0.3 mg protein as determined before denaturation. The glucose background in each sample was determined in a sample without the glucoamylase and subtracted from the sample before calculating the glycogen concentration using the standard curve.

### Microscopy

Saccharomyces cerevisiae strains expressing Hsp104-C-GFP in the WT background, and Hsp104-C-GFP in pin4*_RRMΔ_* background were cultured in 20 mL of SD-Trp medium from OD_600_ of 0.05 to OD_600_ of 0.45. After reaching this optical density, 175 µL of cell suspension was put onto a chambered coverslip with eight individual wells (Thistle Scientific) that has been pre-coated with 10 µL of Concanavalin A (ConA, 5 mg/mL stock) for 10 min at 32°C and subsequently washed three times with SD-Trp medium. Cells were allowed to adhere to the coverslip surface for 20 min at 32°C. Following adhesion, non-adherent cells were removed with three washes using 500 µL of SD-Trp medium. The adherent cells were then imaged at 25 °C using a spinning disc confocal microscope with a Nikon TE2000 microscope base, a modified Yokogawa CSU-10 spinning disk (Visitech) an iXon + Du888 EMCCD camera (Andor) and a 100x/1.45 NA Plan Apo objective (Nikon). For glucose starvation monitoring, the cells were washed three times with 500 µL of glucose-starvation medium (SD-Trp without glucose, supplemented with 2% glycerol and 2% ethanol), and imaged for further 2-hour period. Metamorph software (Molecular Devices) were used to capture images every three min with an exposure time of 150 ms, 488 nm laser set at 35% power and a 0.6 µm step-size over 11 z-sections at three-minute intervals.

### Polysome profiling

WT and *pin4Δ* cells were grown in SD-Trp medium overnight to obtain the saturated cultures the following morning. Thes cultures were then diluted in 100 mL of SD-Trp and grown in synthetic complete (SC) medium from OD_600_ of 0.05 to OD_600_ of 0.45. Cycloheximide was added to the cultures at a final concentration of 0.1 mg/mL for 5 min before harvesting to stabilize ribosomes by blocking them during elongation phase. Cells were subsequently pelleted by centrifugation at 4,600 rpm for 2 min at 4°C, washed once with 20 mL of ice-cold water, snap-frozen in a dry-ice and ethanol bath, and stored at -80°C. Biological duplicates were collected for each condition. For glucose starvation experiments, when reaching the right OD, cells were initially filtered and transferred to 100 mM of SC medium supplemented with 2% glycerol and 2% ethanol instead of glucose, for 16 min. The cells were then harvested as described above.

Following the methodology outlined in Winz et al. (2019)(Winz et al. 2019), cell lysis was performed by resuspending cell pellets in 300 µL of lysis buffer (20 mM HEPES-KOH pH 7.4, 100 mM potassium acetate, 2 mM magnesium acetate and EDTA-free protease inhibitors).

The suspension was transferred to screw-cap tubes containing 200 µL of zirconia beads prepared on ice and subjected to four cycles of bead beating using a Fast Prep-24 machine (MP Biomedicals) set at 6m/s for 40 seconds with a pre-cooled cool adapter. For each of the beating, 8g of dry ice was used to keep the samples cold, but not freezing. An additional 200 µL of lysis buffer was added post-bead beating, and the lysate was cleared twice for 15 min at maximum speed at 4°C. RNA concentration was measured on a Nanodrop spectrophotometer.

300 µg of RNA was loaded onto 10%-45% sucrose gradients made in 1x gradient buffer (10 mM Tris-acetate pH 7.4, 70 mM ammonium acetate, 2 mM magnesium acetate). Gradients were first made using a Gradient Master (BioComp) set to the “Short; SW40; Suc10-45%” program and refrigerated for 2 hours. After loading the RNA on the gradients, the gradients were centrifuged at 38,000 rpm for 2.5 hours at 4°C in SW40-Ti rotors using an Optima XPN-100 Ultracentrifuge (Beckmann Coulter) with acceleration set at 1 and deceleration at 9. Post-centrifugation, gradients were fractionated into 60 fractions of 200 µL each using a Piston Gradient Fractionator (BioComp), equipped with a TRIAX flow cell (BioComp) for UV profiling and an FC203B fraction collector (Gilson).

### Metabolomics

The experiment was performed as described (Bexley et al. 2025). Briefly, cells were grown in SD media, harvested by filtration and either lysed immediately or 10 ml shift media was applied directly to the filter and incubated for the time indicated. Cells were lysed by placing the filter in a petri dish containing 4 ml 12% Trichloroacetic acid (w/v) in 15 mM MgCl_2_ snap frozen in liquid nitrogen. For NTP quantification, lysates were analyzed by HPLC UV as described (Jia et al. 2015). UV chromatograms for experimental samples were accompanied by NTP, NDP and AMP standards. Intracellular nucleotide concentrations were estimated as described (Sabouri et al. 2008) and expressed relative to the value for the unperturbed wildtype in SD.

## QUANTIFICATION AND STATISTICAL ANALYSIS

### Protein domain analysis of Pin4

Regions within Pin4 matching protein domain families were identified using the NCBI Conserved Domains database, which revealed predicted RNA recognition motif (RRM) and R3H domains. Domain alignments and phylogenetics trees were constructed based on the information available on the same platform. 3D structure prediction for Pin4 was performed using AlphaFold3.

AlphaFold3 modelling was carried out with two copies of the full-length sequences of Pin4 from *S. cerevisiae* (Abramson et al. 2024). After inspection, a sequence encompassing the R3H was also modelled, also with two copies (residues 273-373; R3H designated as residues 306-367). The predicted aligned error was visualized using PAE viewer (Elfmann and Stülke 2023) and figures were generated using Pymol.

### reCRAC sequence data analysis

Sequencing of reCRAC libraries was done on the Illumina MiniSeq system, using the High Output Reagent Cartridge. Bioinformatic analysis of reCRAC followed the protocol described by Ristová et al., 2024. Initially, reads were demultiplexed using pyBarcodeFilter.py based on uniquely barcoded 5’ linkers. Quality filtering and 3’ adapter trimming were done using Flexbar with specified settings (--removal-tags -at RIGHT -ao 4 -u 3 -m 7 -n 16 -t), and targeting the adapter sequence TGGAATTCTCGGGTGCCAAGGC. PCR duplicates were removed using pyFastqDuplicateRemover.py, which collapses reads with identical three random nucleotide sequences present in the 5’ linker. These collapsed FASTA reads were then aligned to the Saccharomyces cerevisiae reference genome (SGD v64) using Novoalign (V2.07.00, Novocraft) with Ensembl annotations (EF4.74), applying the Random -r method. Read quantification was executed with pyReadCounters.py, using the Ensembl GTF version EF4.74, which includes mRNA annotations with UTRs, as well as different non-coding RNA categories such as CUTs and SUTs.

The output of pyReadCounters.py is a read count file (hittable). One pseudocount was added to each transcript to improve quantification. The reads were RPM normalized due to a very specific Pin4 binding peak and the normalized data were plotted against RPKM normalized RNAseq data. For the distribution of binding, the ratio between 5’UTR, CDS, 3’ UTR over the genomic region was calculated, mean value from the three replicates was taken with SD.

Most analysis were done on transcripts that showed the strongest Pin4 binding, for these an arbitrary cut off of at least 0.3% of the library was made from the raw reads. Protein binding sites were visualized using the Integrative Genomics Viewere (IGV). For this purpose, reads aligned with Novoalign were converted from SAM to BAM format, which were then sorted and indexed using samtools. Read normalization was achieved by calculating a scaling factor relative to the library size, and coverage was determined using bedtools for each strand, producing scaled bedgraph files for loading onto IGV. The pyPileup.py tool was utilized to illustrate the distribution of reads with number of deletions and substitutions, which represent the sites of protein-RNA crosslinks, along the TPI1, GPM1 and ALT2 genes. The motif sequence was found by taking the targets that make up 0.3% of the library, identified the highest deletion peak and taking 10 nt from upstream and downstream from this peak and searched for enriched motifs using MEME. 2D plotter was used to generate a metagene plot of reads aligned to their 3’ end. Scatter plots and bar graphs were created using GraphPad Prism 9. Gene ontology representations were generated via the Proteomaps website. Bar graphs display the median values from three experimental replicates, with error bars representing the standard deviation (SD).

To evaluate transcript abundance across function gene classes in reCRAC and RNAseq datasets, we performed comparative analysis using selected gene sets of strong Pin4 targets (≥ 0.3% of the total library and ≥1.5-fold enrichment in ≥2 replicates) corresponding to glycolysis, amino acid metabolism, mitochondrial metabolism, and translation. Gene assignments were based on GO annotations and literature search. Transcript abundance values were obtained from three biological replicates of Pin4 reCRAC and RNAseq datasets, under both glucose and stress conditions (16 minutes glucose starvation). For each gene, the mean read count (X) was calculated across the three replicates for each condition. A global mean (x) and standard deviation (SD) were then computed across the four condition specific means for that gene. Each condition specific mean value was then normalized using the z-score formula representing the relative abundance of each transcript:

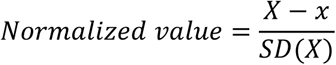

Statistical comparisons were made using Mann-Whitney statistical test.

### RNA-seq analysis

The paired-end Illumina RNA-seq data were first pre-processed using Flexbar to trim low quality bases and adapters from raw reads. The specific parameters set for Flexbar included quality filtering of tail bases with a quality threshold of 30 (Sanger scale), adapter removal with a minimum overlap of four bases, and discarding of reads shorter than 18 bases after trimming (-q TAIL -qf sanger -qt 30 -aa TruSeq -ao 4 -m 18 -n 16). The cleaned reads were then aligned to the *S. cerevisiae* transcriptome using Salmon (Patro et al. 2017) to quantify transcript abundances. For this, the genome GTF file and FASTA files for genes, cDNAs, and ncRNAs were downloaded from Ensembl (version GCF_000146045.2_R64-1-1). These files were used with Salmon’s generateDecoyTranscriptome.sh script to create a hybrid FASTA containing decoy genome sequences concatenated with the transcriptome. Indices for this decoy-aware transcriptome were built using the Salmon indexer. The salmon quant command was then employed to quantify the processed reads against this index. The resulting data were imported into R and processed with the edgeR package. Reads were normalized using the Trimmed Mean of M-values (TMM) method (Robinson and Oshlack 2010). Data filtering was performed using the ‘filterByExp’ function in edgeR, applying a count per million (CPM) threshold with a minimum count per sample of 15 and a minimum total count of 30.

The experimental design was modelled using the ‘model.matrix’ function in R, which accounted for both sample variations and replicates. Dispersion estimates were obtained using ‘estimateDisp’, and generalized linear models (GLM) were fit to the data using the ‘glmQLFit’ function employing a quasi-likelihood approach. Differential expression analysis was subsequently performed using ‘glmQLFTest’. Visualisation of differentially expressed genes, heatmaps, volcano plots, was performed using ggplot2, with significant genes being identified based on a fold change threshold of 2 and a false discovery rate (FDR) of 0.05.

PCA plots were made using the log transformed CPM normalized data.

For comparative analysis with reCRAC data, RNAseq datasets of the WT under normal and glucose starvation conditions were mapped using Novoalign to generate SAM files. These files were then processed to produce bedgraph files as described in the **reCRAC sequence data analysis** section. Visualization of these data was facilitated using the Integrative Genomics Viewer.

### qPCRs

Each sample was measured in a technical and biological triplicate and a median was taken for the results. The relative expression levels were calculated using the ΔΔCt method with a housekeeping gene (ALG9) as the reference. Data were normalized to the control condition and expressed as fold changes. Bar graphs show the average of three replicates and error bars show SD or SEM. Detailed information is provided in the Fig. captions. GraphPad Prism 9 software was used for the statistical analyses.

### Microscopy Image Processing and Analysis

Raw microscopy pictures were processed using ImageJ software (Fiji; NIH). Displayed images represent maximum projections of 11 z-sections with a 0.6 µm step-size. To quantitatively assess the presence of Hsp104 protein granules, cells were categorized as either containing or lacking visible granules at each time point. The percentage of cells with at least one granule was calculated and graphically represented using GraphPad Prism 9 software together with the statistical test. Figure 1B was made using BioRender.

## ACKNOWLEDGEMENTS

We thank Richard Clark from Clinical Research Facility Edinburgh for sequencing services; Shaun Webb and Ola Helwak for answering all our extensive bioinformatics questions; and Adam Kováč and Ye Dee Tay for their help with the spinning disc confocal microscope and with the analysis and processing of raw data images. We thank Ola Helwak and Sophie Giguere for the critical reading of this manuscript. MR was supported by an EASTBIO PhD fellowship [543KOM/G40389]. KB was supported by Wellcome Trust PhD Studentship [218470]. SS and AC were supported by the Swedish Cancer Society [22 2377 Pj] and Swedish Research Council [2022–00675_VR]. JFS, VS and DT were supported by Wellcome Principal Research Fellowships [109916, 222516]. AGC was supported by a Wellcome Senior Fellowship [200898]. This work was facilitated by funding for the Wellcome Discovery Research Platform for Hidden Cell Biology [226791] and we gratefully acknowledge support from the Microscopy and Bioinformatics cores. Work in the Wellcome Centre for Cell Biology was supported by Centre Core Grants (092076 and 203149).

## SUPPLEMENTARY FIGURE LEGENDS

## Supplementary Tables

Supplementary Table 1:

The list of oligonucleotides used in this study.

Supplementary Table 2:

The list of strains generated in this study.

Supplementary Table 3:

The list of plasmids generated in this study.

Supplementary Table 4:

List of Pin4 reCRAC 3’ UTR and CDS binding mRNAs recovered in reCRAC following glucose withdrawal for 16 min.

Supplementary Table 5:

List of mRNAs recovered in Pin4 reCRAC.

Supplementary Table 6:

Comparison of mRNA recovery in Pin4 reCRAC in glucose medium with RNA sequencing.

Supplementary Table 7:

Differentially expressed mRNAs +/- Pin4 on glucose medium.

Supplementary Table 8:

Differentially expressed mRNAs +/- Pin4 after glucose withdrawal for 16 min.

---

**Supplementary Figure 1:**
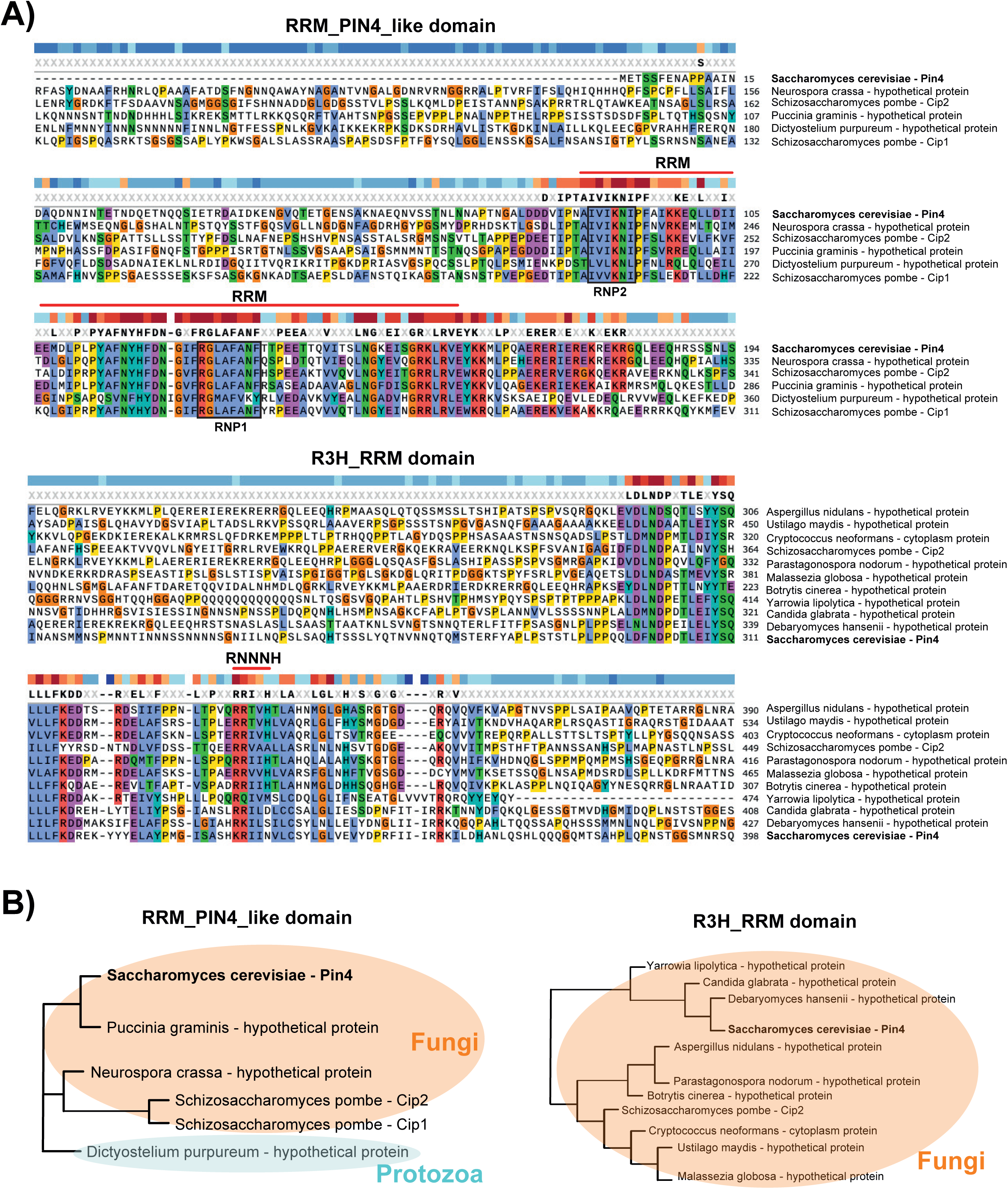
Domain structure of Pin4 A) Multiple sequence alignment of the conserved domains of Pin4: RRM_PIN4_like and R3H_RRM domains across various species. The conserved residues are colored, and the beginning and end of both the RRM and R3H domains are annotated. RNP1 and RNP2 motifs within RRM are also annotated. B) Phylogenetic trees depicting the evolutionary relationship of the RRM_PIN4_like and R3H_RRM domain containing proteins. The fungi are highlighted in orange, and the protozoan species in blue.

**Supplementary Figure 2:**
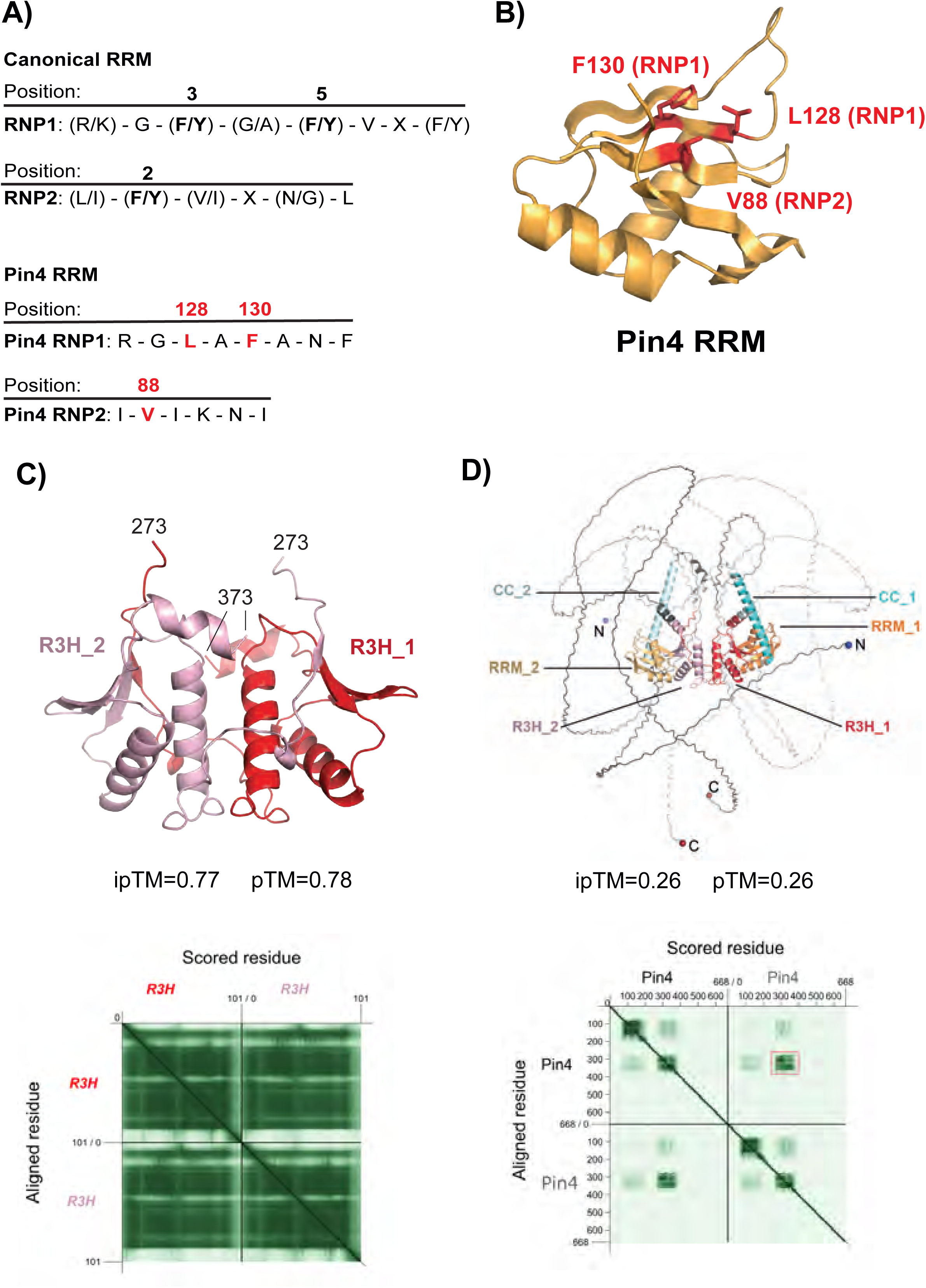
Pin4 RRM domain is non-canonical and R3H may homodimerize A) Comparison of RRM motifs between the canonical RRM and Pin4 RRM. The diagram indicates critical amino acid positions and sequences within RNP1 and RNP2 of the canonical RRM, and alterations in Pin4. B) Modelled 3D structure of the Pin4 RRM domain, Non-consensus amino acids, L128, F130 (RNP1), and V88 (RNP2) are highlighted in red. C) AlphaFold3 prediction of the R3H domain dimer from Pin4. The domains are colored in red or pink and the residues at the N and C terminal positions of this fragment are marked with residue numbers. PAE matrix for models with the R3H domains alone modelled as a dimer. D) AlphaFold3 prediction of dimeric Pin4, with RRM domains colored light and bright orange, coiled coil (CC) regions colored light and bright cyan, and R3H domains colored red and pink. Note that the R3H domain modeled here extends beyond the annotation in Figure 1A (residues 273-373). The N and C termini are marked with blue and dark red spheres, respectively. Probability scores (ipTM and pTM) are marked. Predicted aligned error matrix (PAE) for full-length Pin4 modelled as a dimer. The area of the matrix corresponding to the predicted R3H interaction is marked with a salmon-colored box.

**Supplementary Figure 3:**
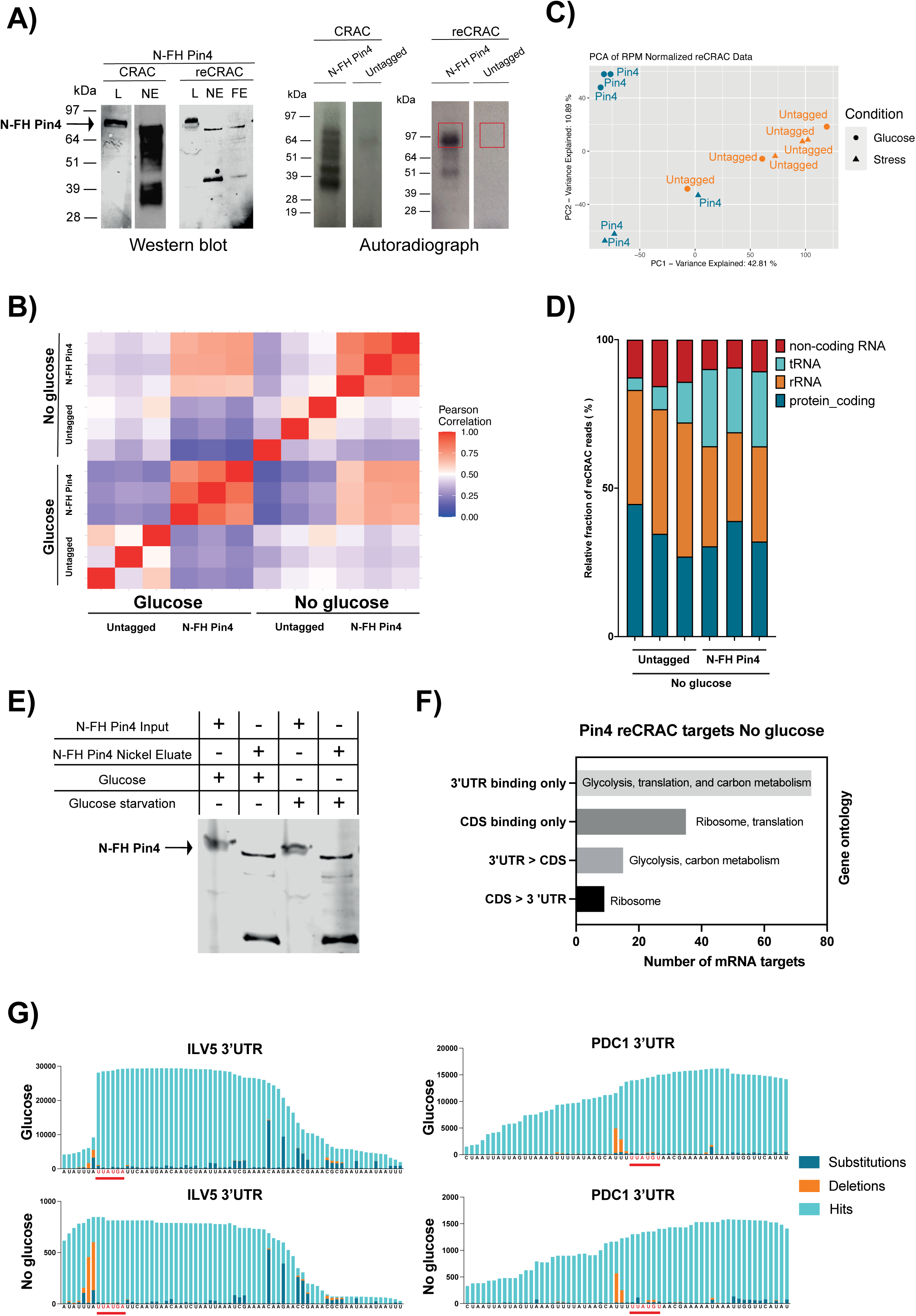
Analysis of Pin4-RNA interactions with reCRAC A) Western blot (left) and autoradiograph (right) analysis showing the protein expression and protein-RNA crosslinking of N-terminal FLAG-Ala_4_-His_8_ tagged (N-FH) Pin4 during CRAC and reCRAC experiments. L, lysate; NE, Ni-NTA eluate; FE, FLAG eluate. The untagged control expresses endogenous Pin4 without the tag. The red boxes indicate the regions excised for further steps. B) Heatmap of Pearson correlation coefficients for N-FH-Pin4 and untagged reCRAC samples performed across glucose and glucose depletion conditions performed in triplicate. The heatmap colors range from blue (low correlation) to red (high correlation). C) Principal component analysis (PCA) of RPM normalized reCRAC data for all RNA classes across different strains and conditions. D) Bar chart displaying the relative fraction of reCRAC reads recovered in No glucose (16 min in Gly/EtOH) from N-FH-Pin4 and untagged strains. The experiment was performed in triplicate for each condition. E) The western blot analysis of the N-FH (FLAG-4xAla-8xHis) tagged Pin4 protein in the input and after the elution from Nickel purification step in glucose and glucose starvation conditions. F) Bar chart of N-FH-Pin4 reCRAC targets following glucose depletion. The data are grouped based on 3’UTR and CDS association (y-axis) and the number of mRNA targets is plotted on the x-axis. Gene ontology enrichment is given for each group. G) The zoomed-in view of reCRAC pileups of the 3’ UTR of Pin4 targets, *ILV5*, and *PDC1* in glucose rich and glucose-poor conditions. The motif sequence UA/UAUGA/U is marked in red. The deletion peaks (orange) represent the sites of crosslinking between RNA and protein

**Supplementary Figure 4:**
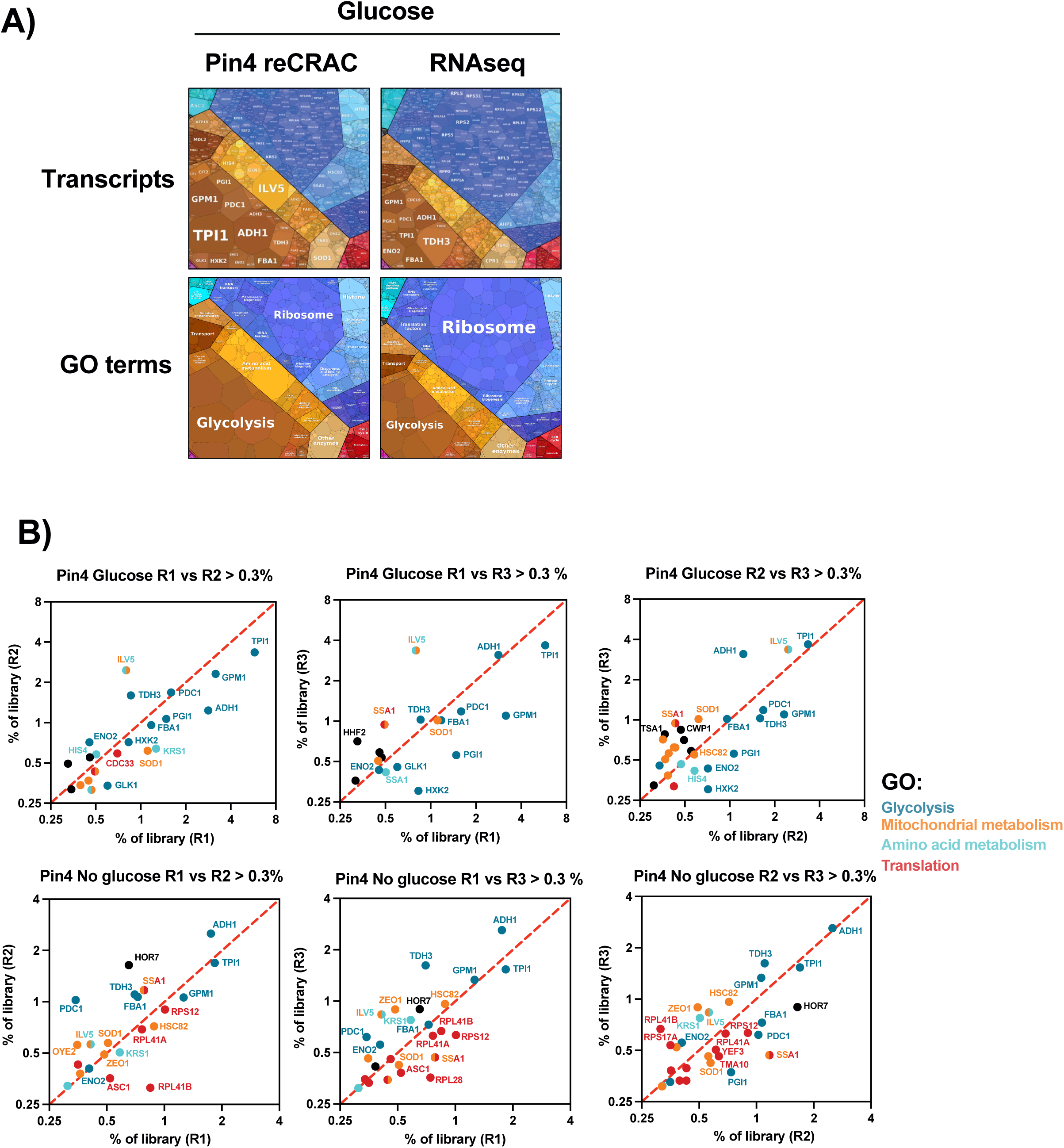
Pin4 targets are enriched for metabolic and pro-growth mRNAs A) GO term enrichment across all Pin4 targets in glucose condition (n=3), and RNAseq data in glucose condition (n=3). Each bubble represents a transcript, and the size of the bubble determines the enrichment of that transcript in a library. Figure prepared using the Proteomaps tool (Liebermeister et al. 2014). B) Scatterplots showing the top Pin4 target mRNAs in glucose and glucose depletion (16 min Gly/EtOH) conditions that represented at least 0.3% of the Pin4 reCRAC library comparing all the replicates. The mRNAs are enriched for four GO term categories: glycolysis, mitochondrial, amino acid metabolism, and translation.

**Supplementary Figure 5:**
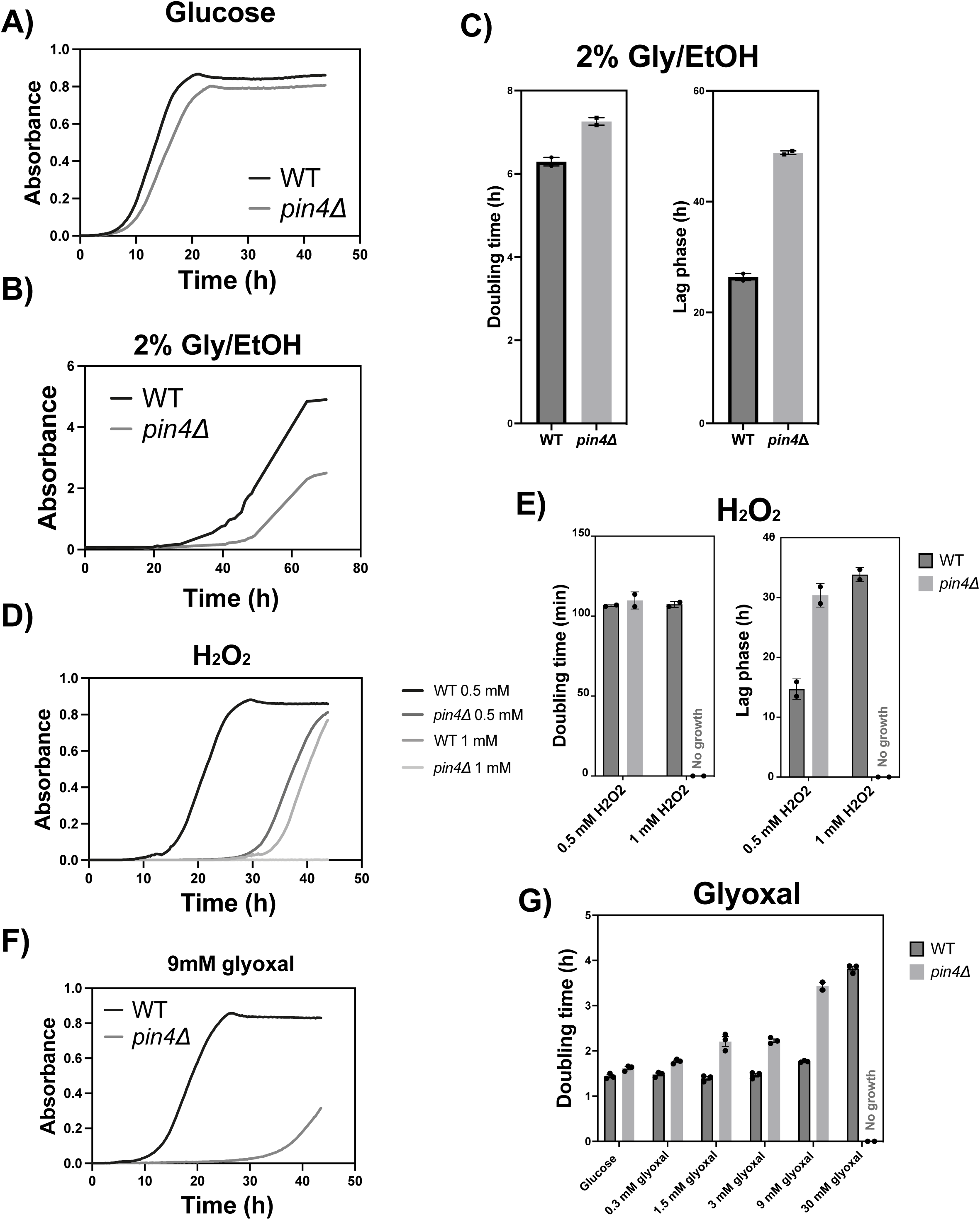
Pin4 deficient cells are sensitive to glucose depletion and mitochondrial stress A) Growth curves of the WT and *pin4Δ* strains in the presence of glucose (n=2, mean value plotted). Absorbance was measured over 45 h to measure growth. B) Growth curves of WT and *pin4Δ* strains during following transfer from glucose medium to medium with Gly/EtOH (n=2, mean value plotted). Absorbance was measured over 70 h. C) Bar chart showing the doubling time and lag phase in hours following shift from glucose to Gly/EtOH. D) Growth curves of WT and *pin4Δ* strain exposed to medium containing varying concentrations of hydrogen peroxide (H_2_O_2_) (n=2, mean value plotted). E) Comparison of doubling time and lag phase of WT and *pin4Δ* strain exposed to 0.5 mM and 1 mM H_2_O_2_. F) Growth curves showing the WT and *pin4Δ* in 9 mM glyoxal (n=2, mean value plotted). G) Doubling times of WT and *pin4Δ* strains in medium supplemented different concentrations of glyoxal (n=2).

**Supplementary Figure 6:**
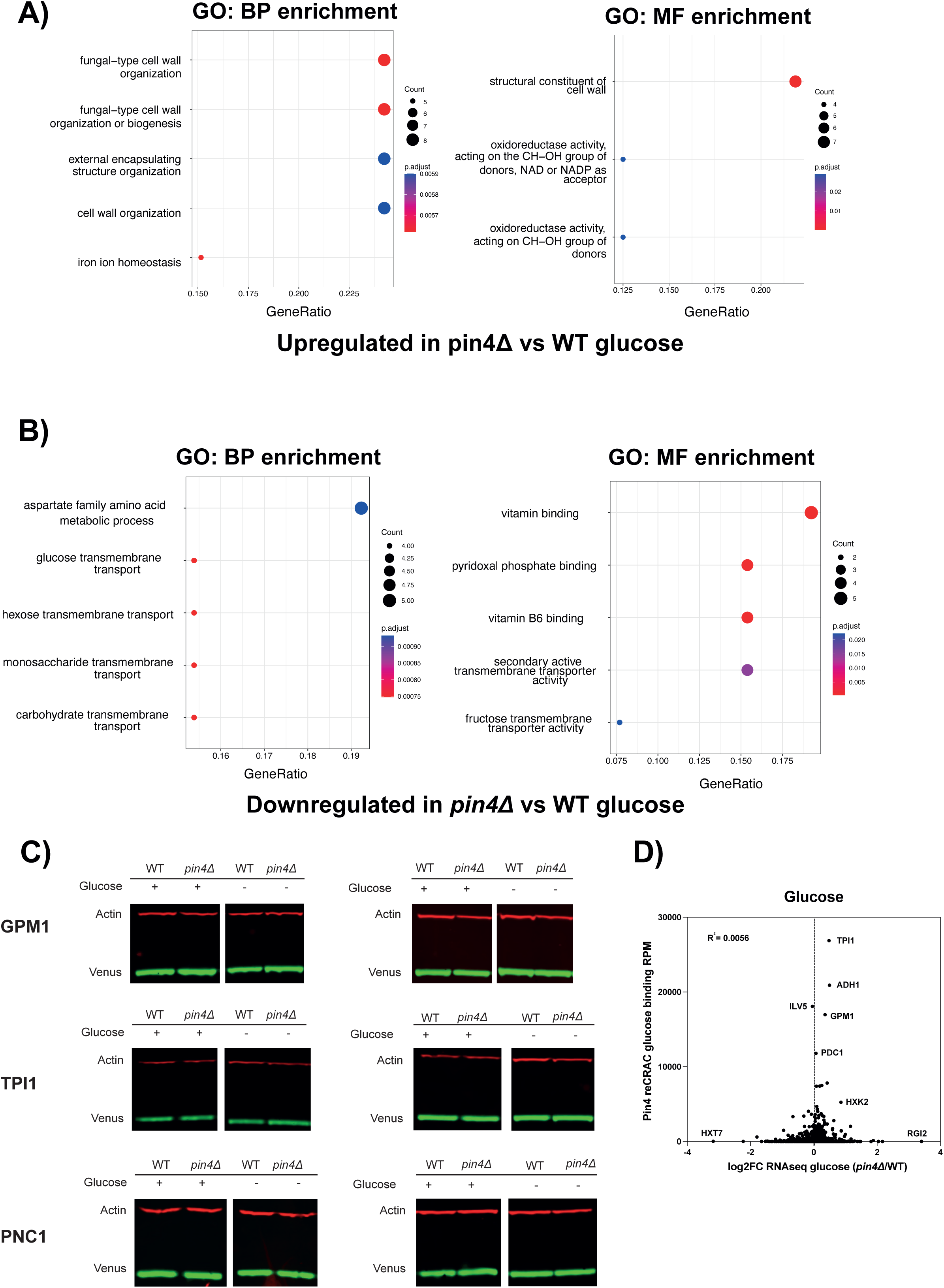
Altered mRNAs and reporters in *pin4Δ* A-B) Bubble charts of GO term enrichment of biological process or molecular function, among mRNAs with increased or decreased abundance in *pin4Δ* relative to the WT in glucose medium. The size of each bubble indicates the gene count, while the color reflects the p-value. Gene ratio is defined as the proportion of differentially expressed genes (DEGs) that are present in a particular gene set, divided by the total number of DEGs. C) Western blots of the Venus reporters with terminator sequences obtained from *GPM1*, *TPI1*, or *PNC1* gene in glucose or glucose depletion (2 hr Gly/EtOH) conditions. The Venus level was normalized to an Actin loading control. D) Scatterplot showing the relationship between RNA-seq log2FC of gene expression in *pin4Δ* compared to WT in glucose and Pin4-RNA binding in this condition. Each point represents a gene. R^2^ value of 0.0056 is shown for the line of best fit.

**Supplementary Figure 7:**
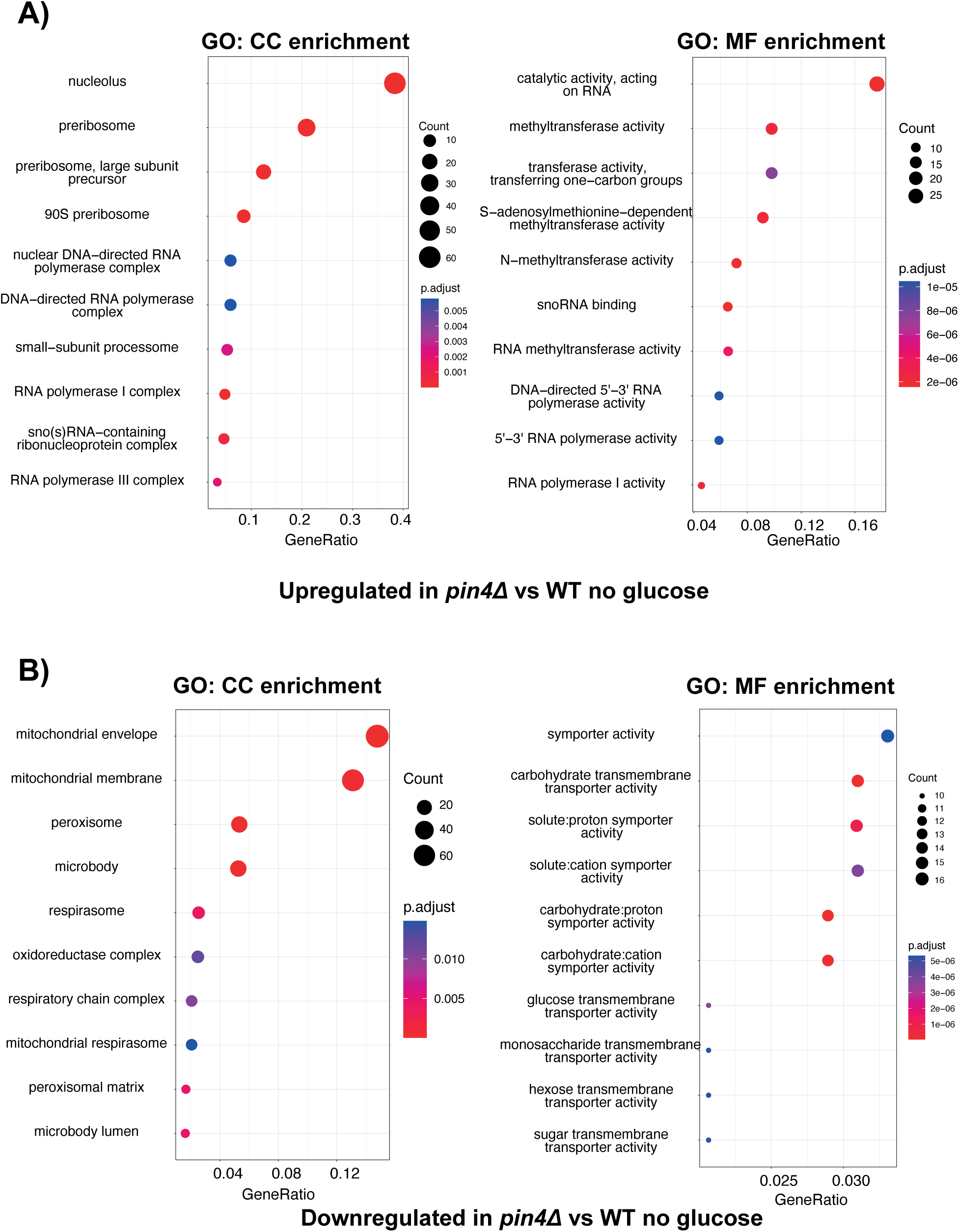
GO enrichment of *pin4Δ* mRNA changes after glucose depletion A-B) Bubble charts of GO term enrichment of cellular component or molecular function among mRNAs with increased or decreased abundance in *pin4Δ* relative to the WT following glucose depletion (16 min Gly/EtOH). The size of each bubble indicates the gene count, while colors reflect the p-value. Gene ratio is defined as the proportion of differentially expressed genes (DEGs) that are present in a particular gene set, divided by the total number of DEGs.

**Supplementary Figure 8:**
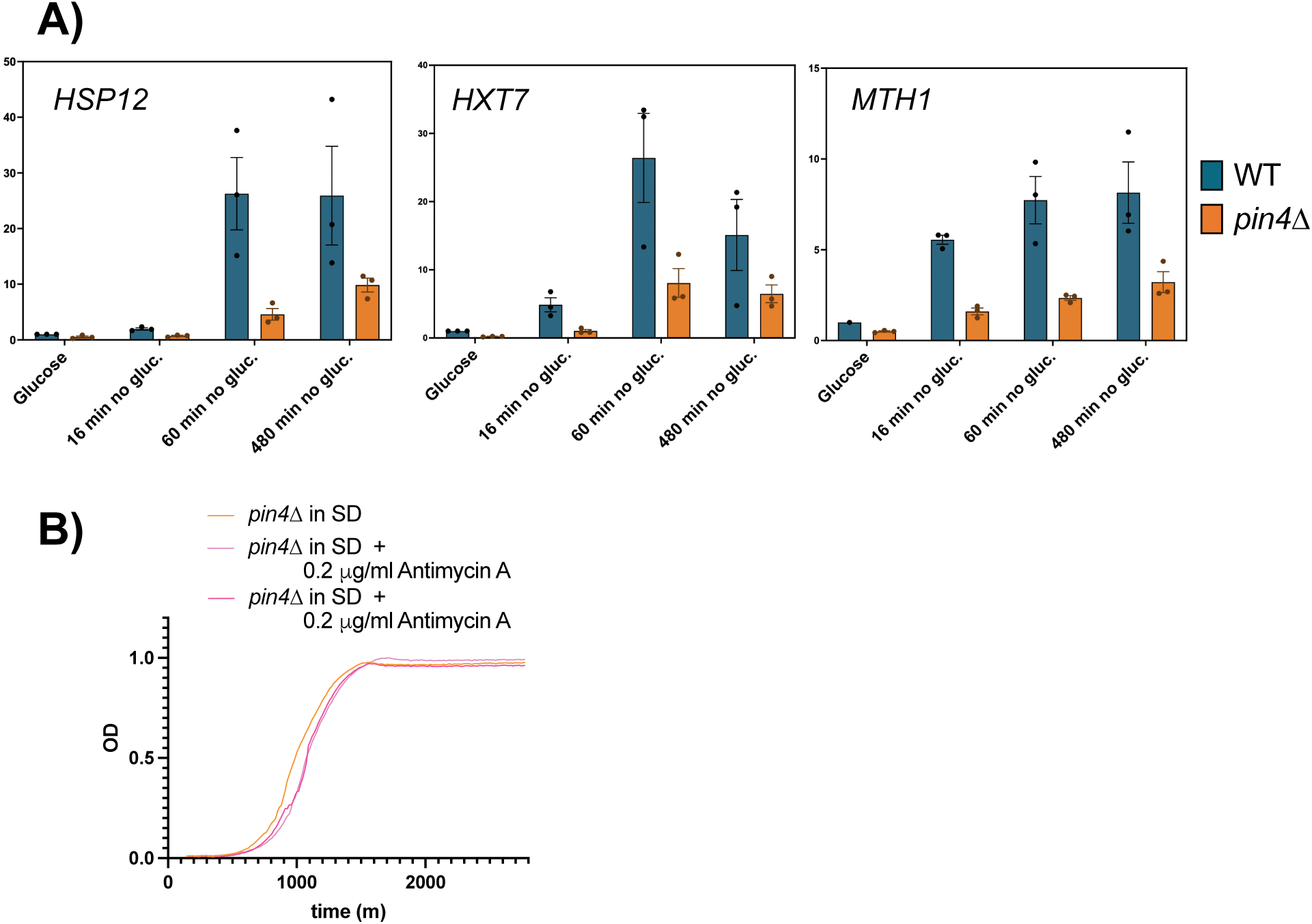
Long-term glucose starvation and AA-treated growth in *pin4Δ* vs WT A) Analyses of gene expression at longer time points. Bar graphs comparing RNA expression levels determined by RT-qPCR normalized to *ALG9*, for WT and *pin4Δ* strains in glucose medium and following glucose depletion for 16, 60 or 480 min (n=3, mean with SEM). Individual replicate values are plotted. B) Growth curves assessing growth in the presence of the mitochondrial inhibitor Antimycin A (AA) in *pin4Δ* cells. Exponential cultures were treated with 0.2 μg/ml, 0.02 μg/ml or no AA and the growth rate was measured by change in optical density (OD) over 48 hours.

**Supplementary Figure 9:**
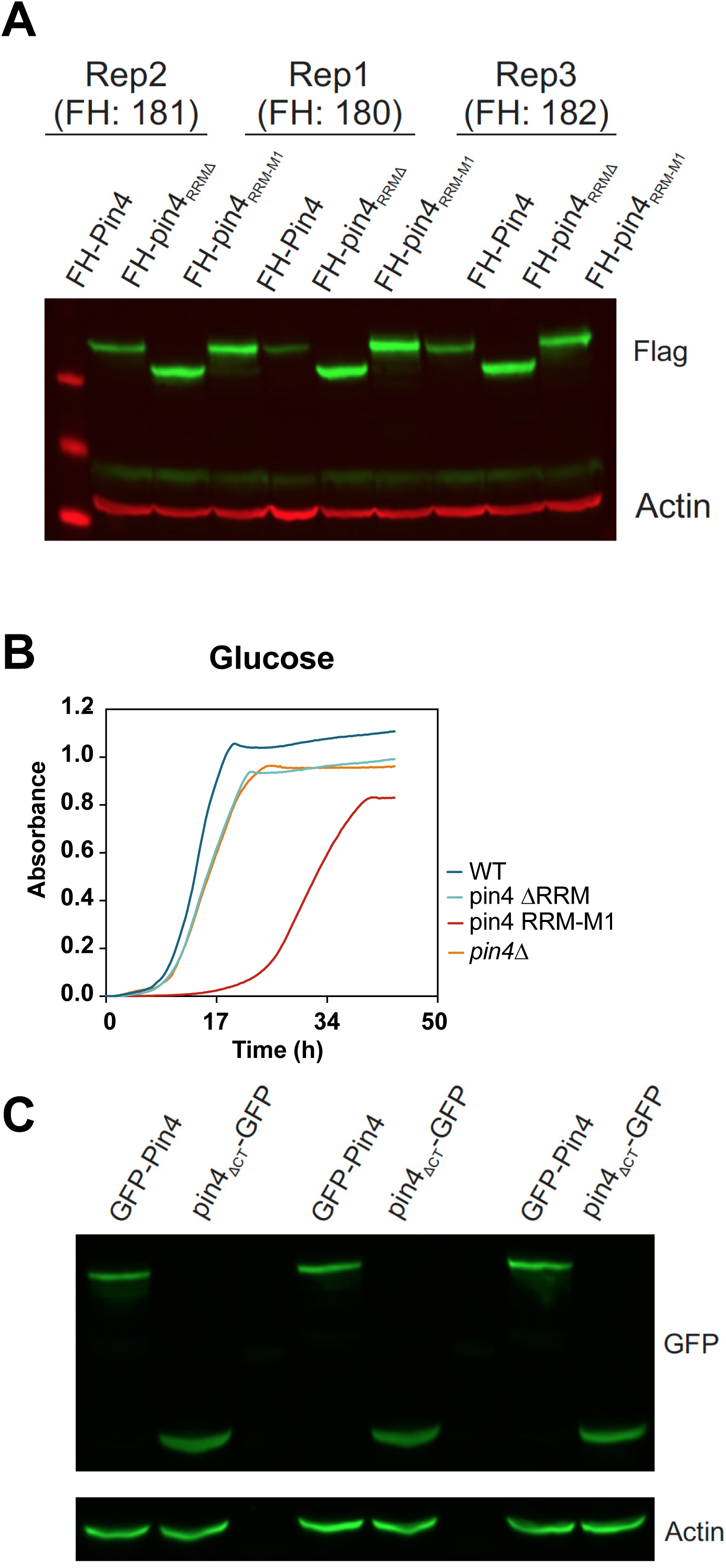
The expression and growth of Pin4 RRM mutants A) Western blot of N-FH tagged Pin4 protein, *pin4 RRMΔ*, and *pin4 RRM-M1* in glucose medium. Three independent sets of strains were analyzed. B) Growth curve of WT, *pin4Δ*, *pin4 RRMΔ*, and *pin4 RRM-M1* in glucose medium. C) Western blot of N-GFP tagged Pin4 protein and Pin4_ΔCT_-GFP in glucose medium

**Supplementary Figure 10:**
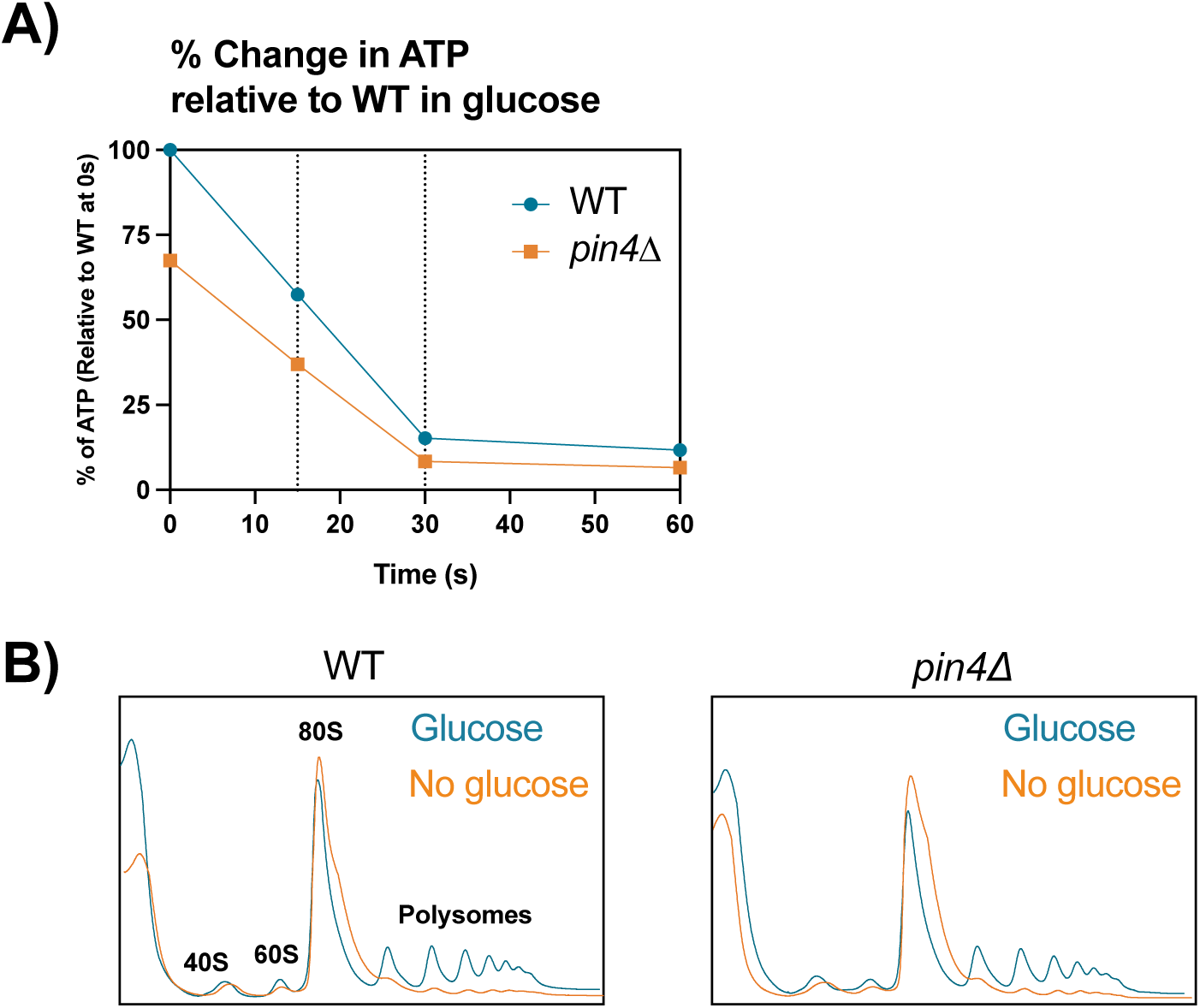
Yeast lacking Pin4 retain the initial stress responses A) Estimated intracellular ATP pool in WT and *pin4Δ strains* following carbon source shift from 2% glucose to 2% glycerol/ 2% ethanol. Mean NTP levels (n=3) quantified at 0 sec,15 sec, 30 sec and 1 min following loss of glucose were converted to percentage change in ATP relative to WT in glucose. B) Polysome profile analysis of WT and *pin4Δ* in glucose (blue) and no glucose (16 min in Gly/EtOH) conditions (orange).

**Supplementary Table 1.**
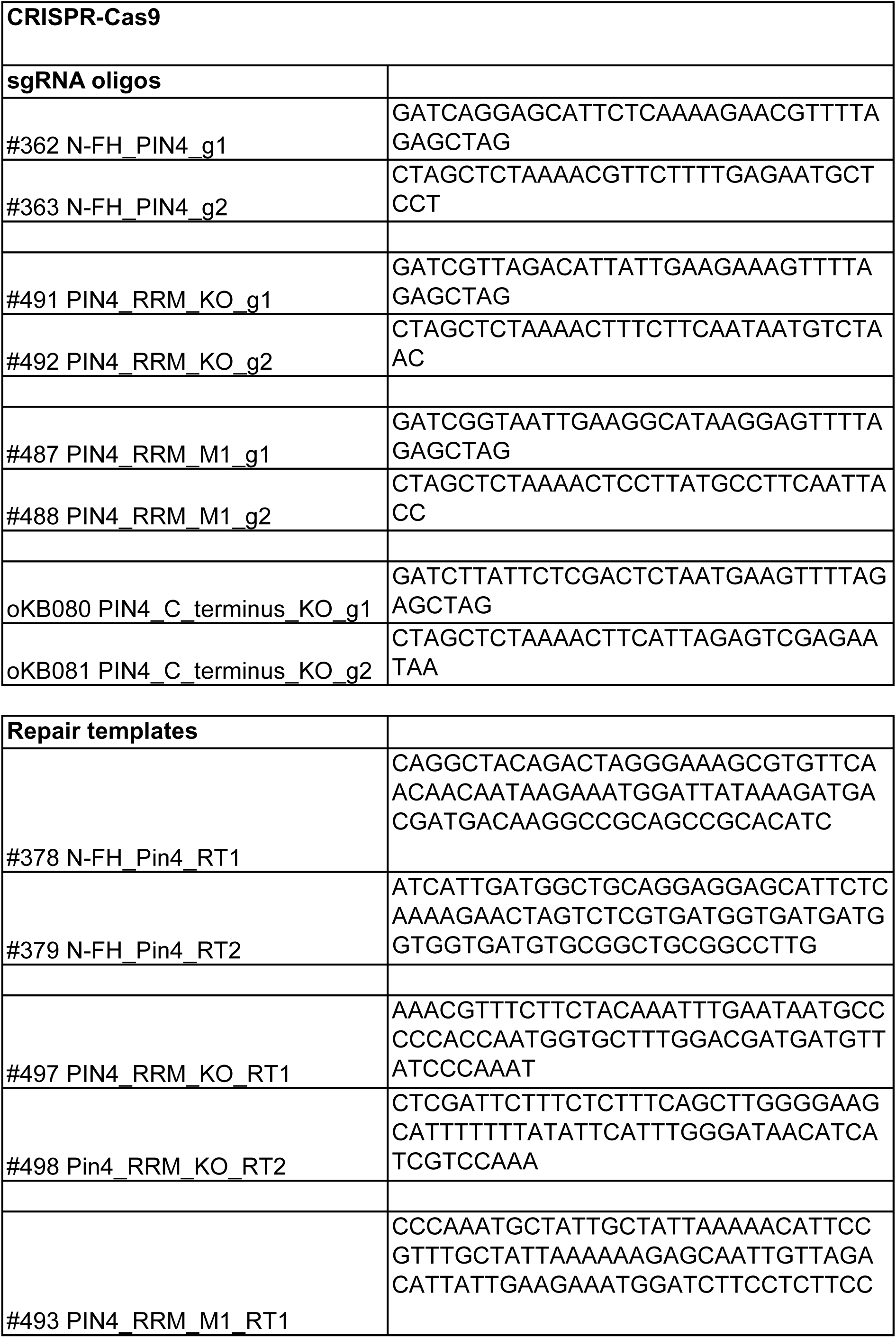
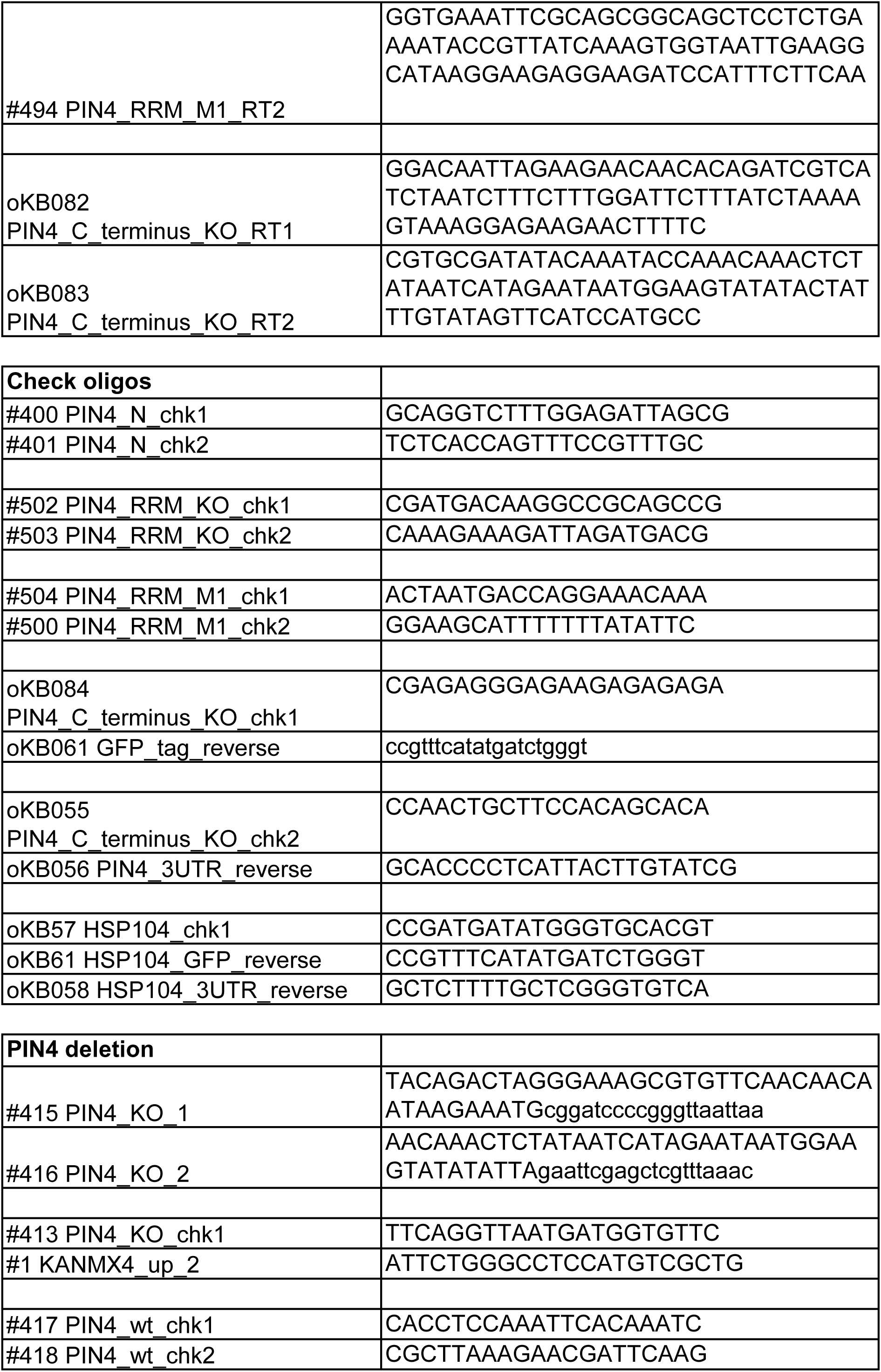
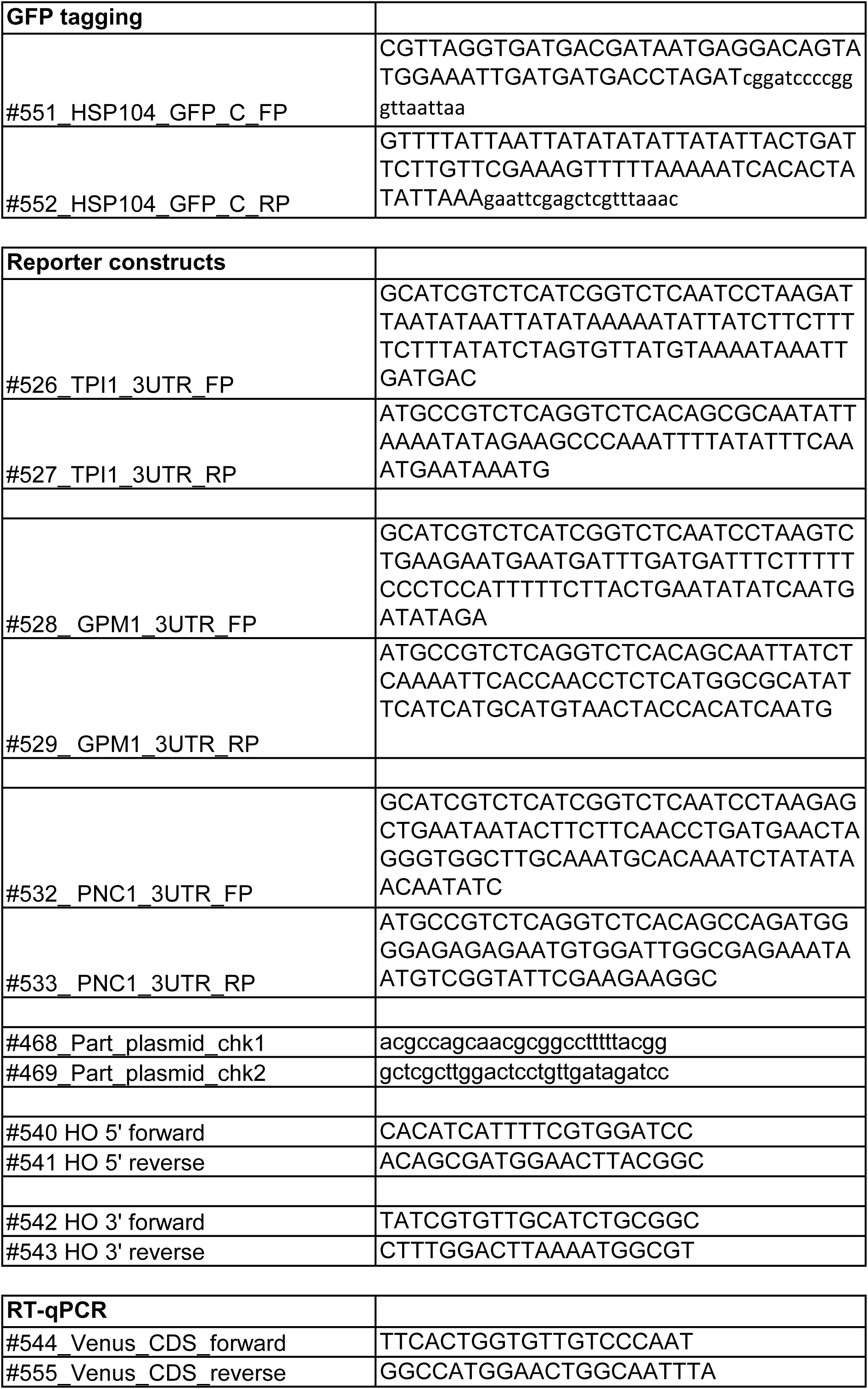
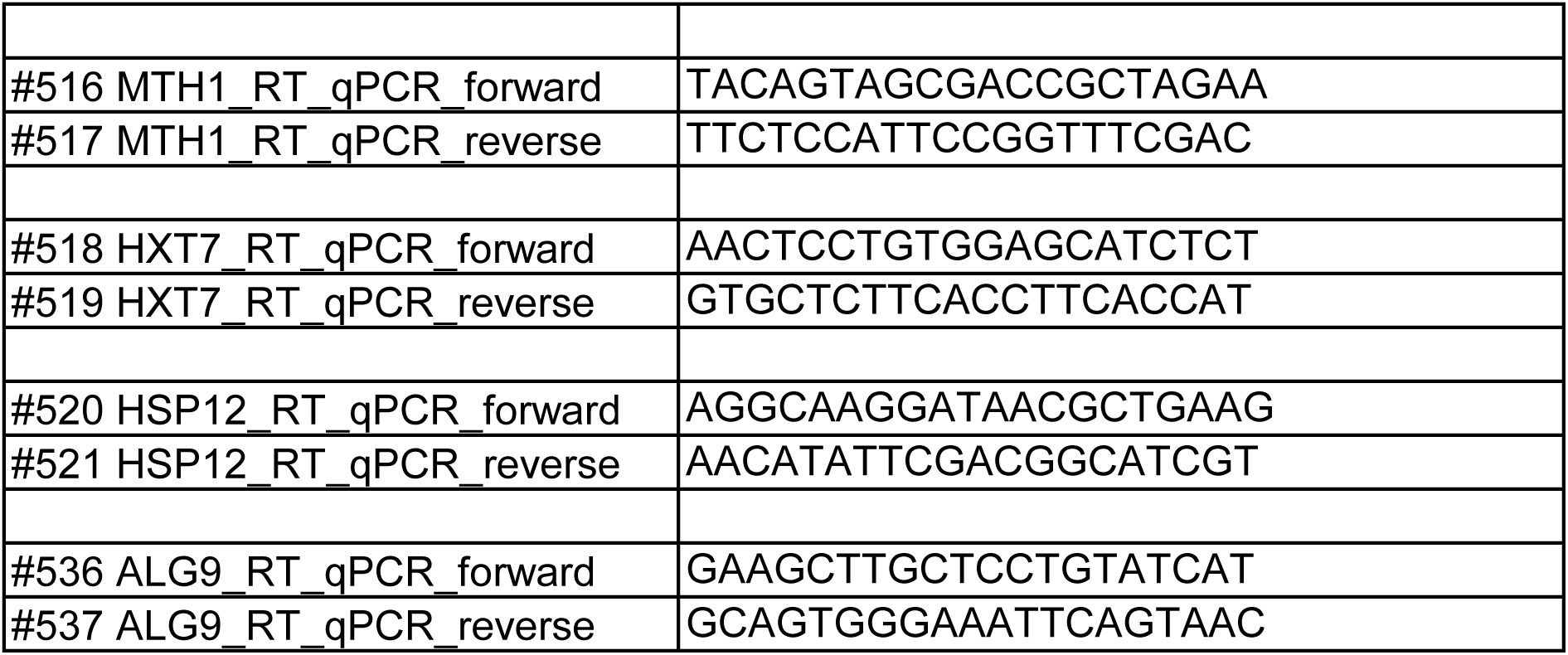
The list of oligonucleotides used in this study.

**Supplementary Table 2.**
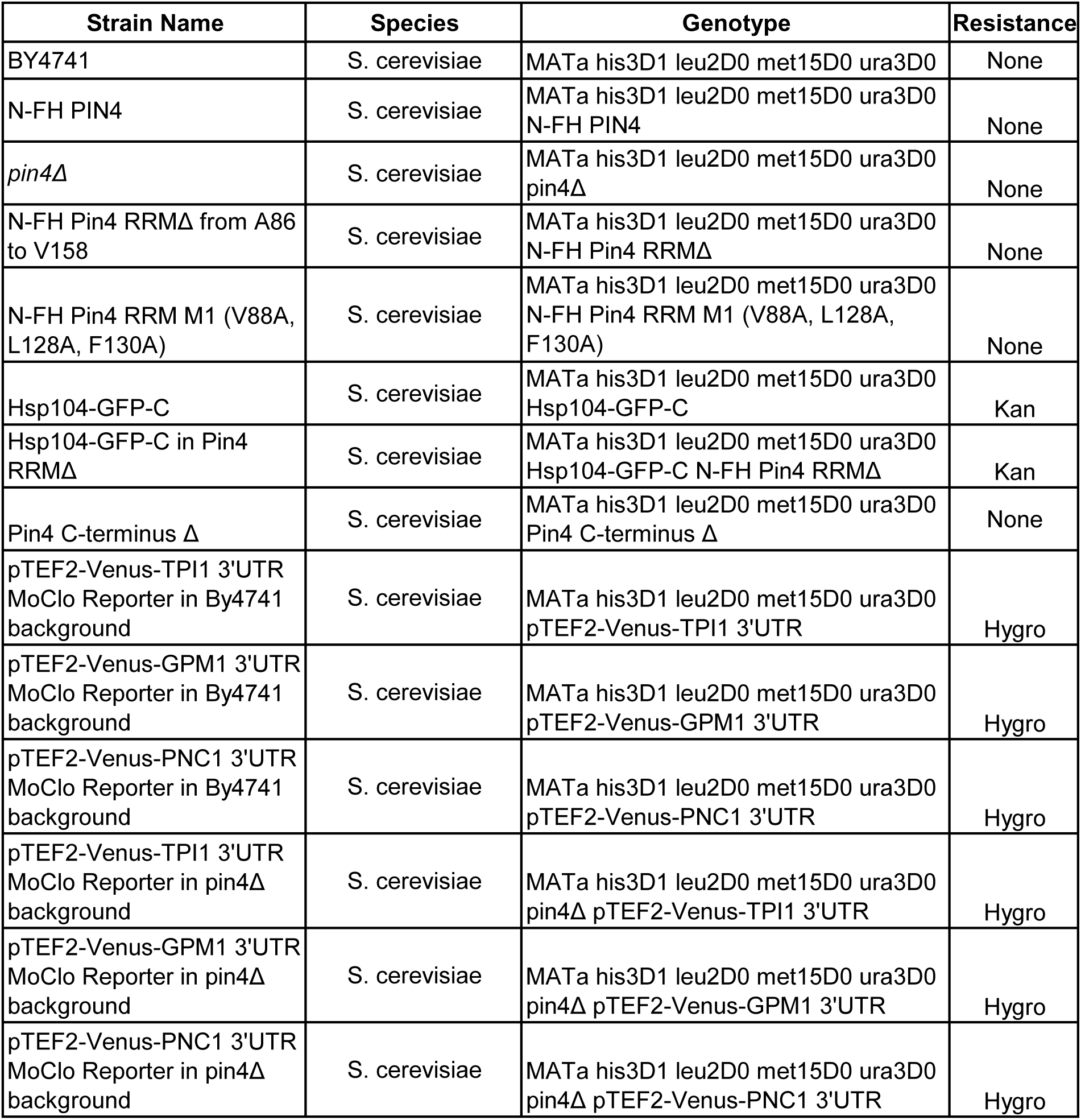
The list of strains generated in this study.

| Strain Name | Species | Genotype | Resistance |
| --- | --- | --- | --- |
| BY4741 | *S. cerevisiae* | MATa his3D1 leu2D0 met15D0 ura3D0 | None |
| N-FH PIN4 | *S. cerevisiae* | MATa his3D1 leu2D0 met15D0 ura3D0<br>N-FH PIN4 | None |
| *pin4*Δ | *S. cerevisiae* | MATa his3D1 leu2D0 met15D0 ura3D0<br>*pin4*Δ | None |
| N-FH Pin4 RRMΔ from A86 to V158 | *S. cerevisiae* | MATa his3D1 leu2D0 met15D0 ura3D0<br>N-FH Pin4 RRMΔ | None |
| N-FH Pin4 RRM M1 (V88A, L128A, F130A) | *S. cerevisiae* | MATa his3D1 leu2D0 met15D0 ura3D0<br>N-FH Pin4 RRM M1 (V88A, L128A, F130A) | None |
| Hsp104-GFP-C | *S. cerevisiae* | MATa his3D1 leu2D0 met15D0 ura3D0<br>Hsp104-GFP-C | Kan |
| Hsp104-GFP-C in Pin4 RRMΔ | *S. cerevisiae* | MATa his3D1 leu2D0 met15D0 ura3D0<br>Hsp104-GFP-C N-FH Pin4 RRMΔ | Kan |
| Pin4 C-terminus Δ | *S. cerevisiae* | MATa his3D1 leu2D0 met15D0 ura3D0<br>Pin4 C-terminus Δ | None |
| pTEF2-Venus-TPI1 3'UTR MoClo Reporter in By4741 background | *S. cerevisiae* | MATa his3D1 leu2D0 met15D0 ura3D0<br>pTEF2-Venus-TPI1 3'UTR | Hygro |
| pTEF2-Venus-GPM1 3'UTR MoClo Reporter in By4741 background | *S. cerevisiae* | MATa his3D1 leu2D0 met15D0 ura3D0<br>pTEF2-Venus-GPM1 3'UTR | Hygro |
| pTEF2-Venus-PNC1 3'UTR MoClo Reporter in By4741 background | *S. cerevisiae* | MATa his3D1 leu2D0 met15D0 ura3D0<br>pTEF2-Venus-PNC1 3'UTR | Hygro |
| pTEF2-Venus-TPI1 3'UTR MoClo Reporter in *pin4*Δ background | *S. cerevisiae* | MATa his3D1 leu2D0 met15D0 ura3D0<br>*pin4*Δ pTEF2-Venus-TPI1 3'UTR | Hygro |
| pTEF2-Venus-GPM1 3'UTR MoClo Reporter in *pin4*Δ background | *S. cerevisiae* | MATa his3D1 leu2D0 met15D0 ura3D0<br>*pin4*Δ pTEF2-Venus-GPM1 3'UTR | Hygro |
| pTEF2-Venus-PNC1 3'UTR MoClo Reporter in *pin4*Δ background | *S. cerevisiae* | MATa his3D1 leu2D0 met15D0 ura3D0<br>*pin4*Δ pTEF2-Venus-PNC1 3'UTR | Hygro |

**Supplementary Table 3.**
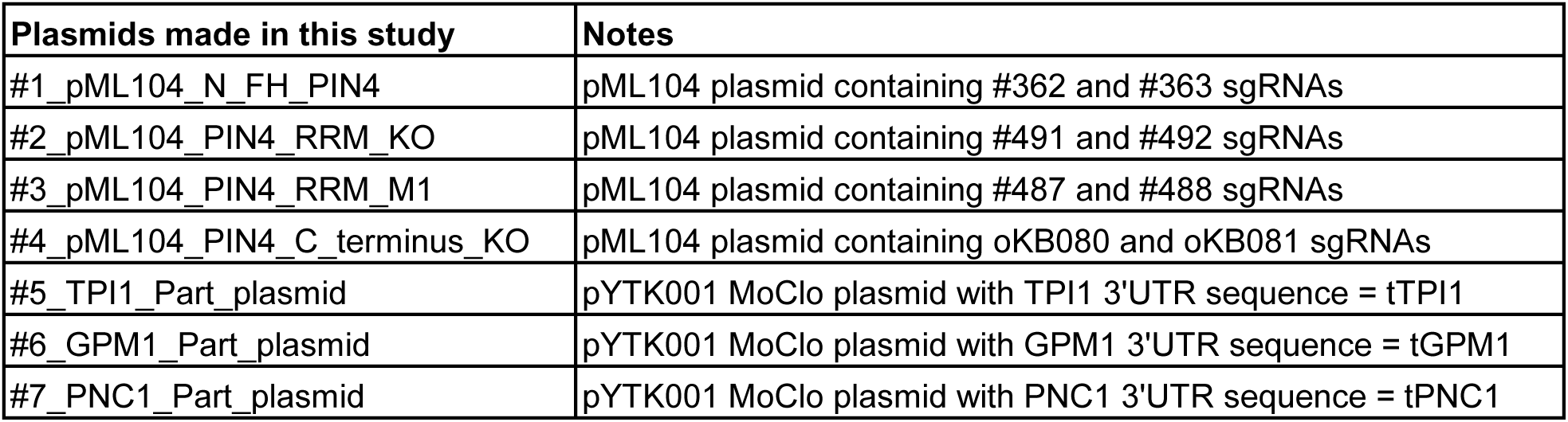
The list of plasmids generated in this study.

**Supplementary Table 4.**
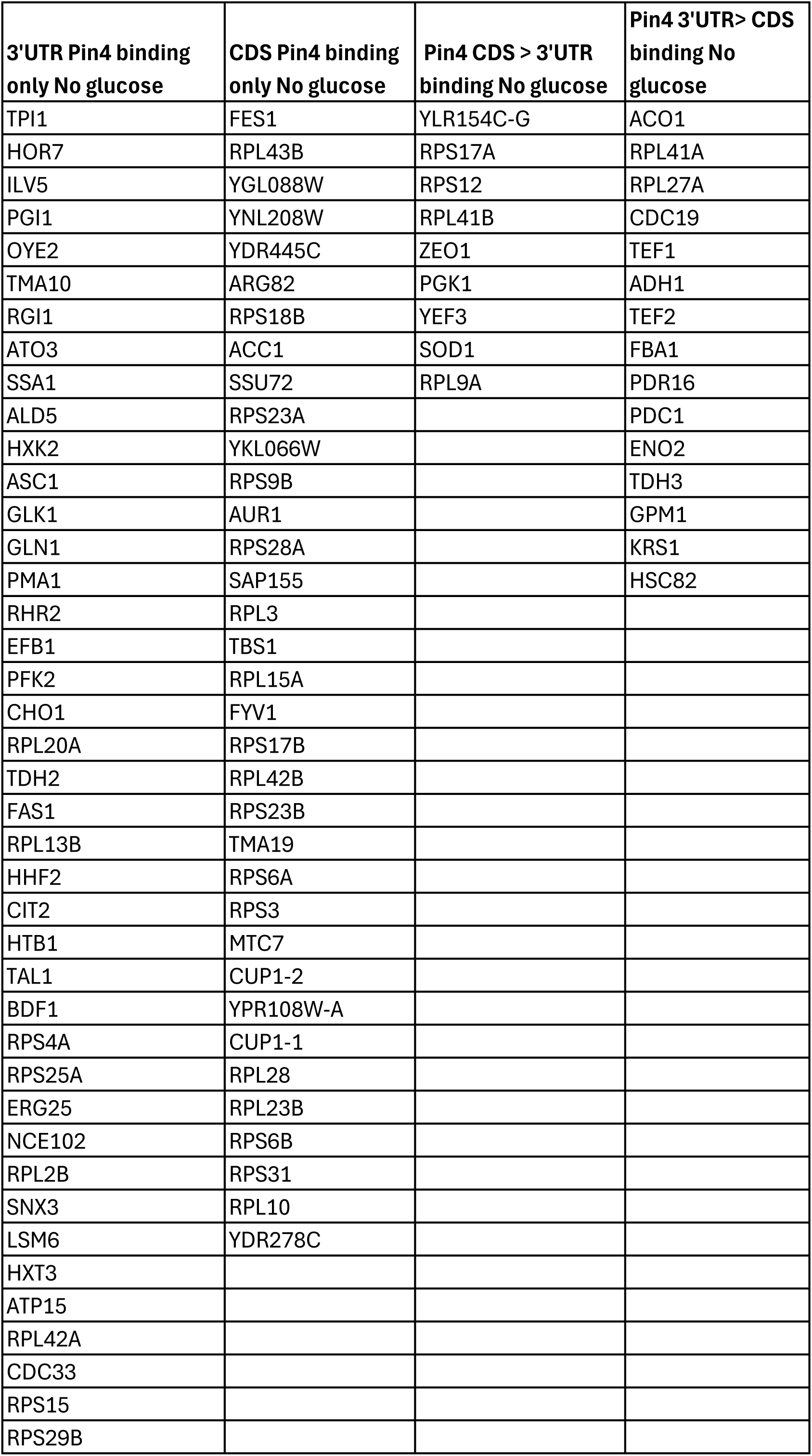
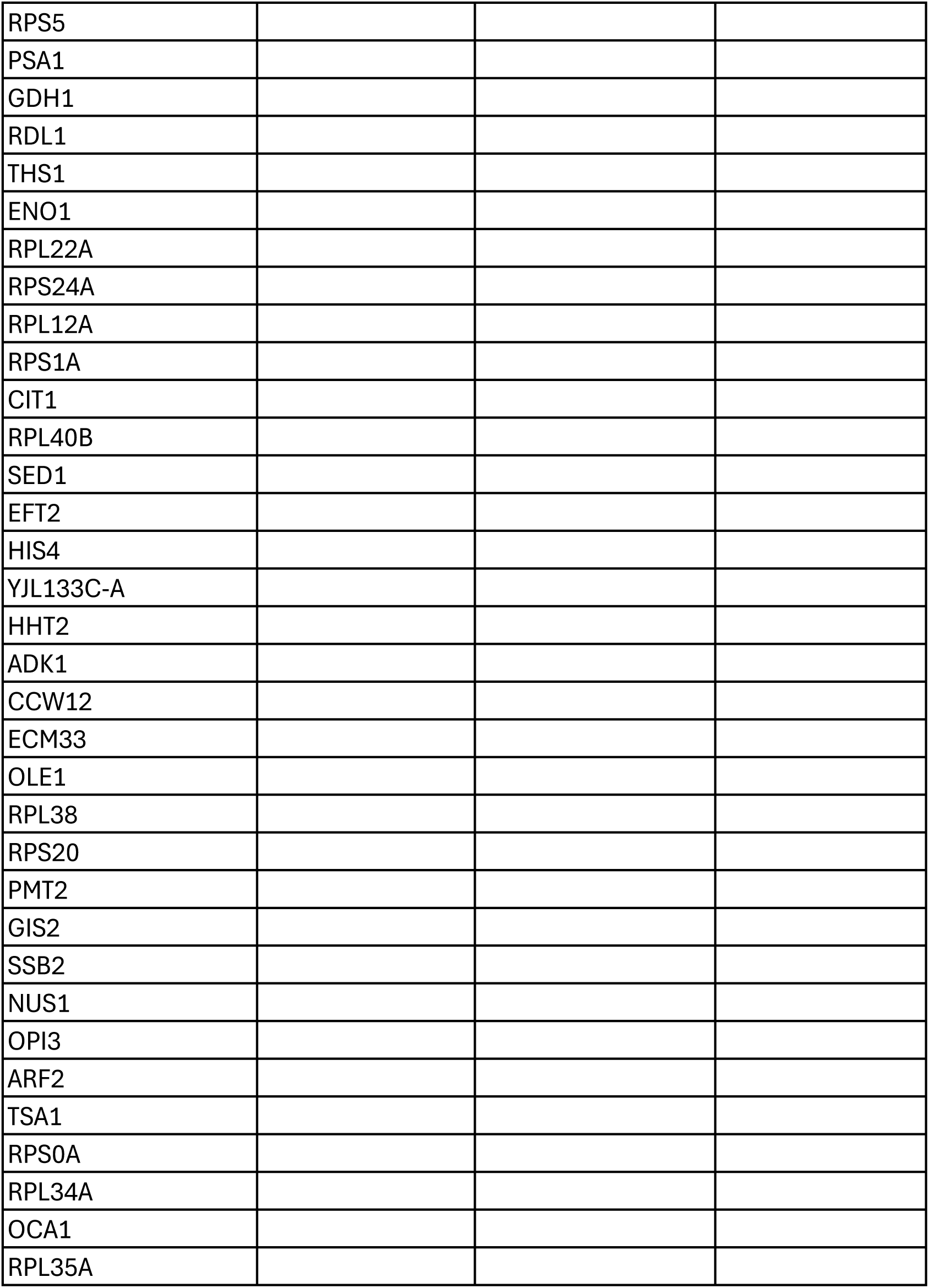
List of Pin4 reCRAC 3’ UTR and CDS binding mRNAs recovered in reCRAC following glucose withdrawal for 16 min.

| **3'UTR Pin4 binding**<br>**only No glucose** | **CDS Pin4 binding**<br>**only No glucose** | **Pin4 CDS > 3'UTR**<br>**binding No glucose** | **Pin4 3'UTR> CDS**<br>**binding No**<br>**glucose** |
| --- | --- | --- | --- |
| TPI1 | FES1 | YLR154C-G | ACO1 |
| HOR7 | RPL43B | RPS17A | RPL41A |
| ILV5 | YGL088W | RPS12 | RPL27A |
| PGI1 | YNL208W | RPL41B | CDC19 |
| OYE2 | YDR445C | ZEO1 | TEF1 |
| TMA10 | ARG82 | PGK1 | ADH1 |
| RGI1 | RPS18B | YEF3 | TEF2 |
| ATO3 | ACC1 | SOD1 | FBA1 |
| SSA1 | SSU72 | RPL9A | PDR16 |
| ALD5 | RPS23A |  | PDC1 |
| HXK2 | YKL066W |  | ENO2 |
| ASC1 | RPS9B |  | TDH3 |
| GLK1 | AUR1 |  | GPM1 |
| GLN1 | RPS28A |  | KRS1 |
| PMA1 | SAP155 |  | HSC82 |
| RHR2 | RPL3 |  |  |
| EFB1 | TBS1 |  |  |
| PFK2 | RPL15A |  |  |
| CHO1 | FYV1 |  |  |
| RPL20A | RPS17B |  |  |
| TDH2 | RPL42B |  |  |
| FAS1 | RPS23B |  |  |
| RPL13B | TMA19 |  |  |
| HHF2 | RPS6A |  |  |
| CIT2 | RPS3 |  |  |
| HTB1 | MTC7 |  |  |
| TAL1 | CUP1-2 |  |  |
| BDF1 | YPR108W-A |  |  |
| RPS4A | CUP1-1 |  |  |
| RPS25A | RPL28 |  |  |
| ERG25 | RPL23B |  |  |
| NCE102 | RPS6B |  |  |
| RPL2B | RPS31 |  |  |
| SNX3 | RPL10 |  |  |
| LSM6 | YDR278C |  |  |
| HXT3 |  |  |  |
| ATP15 |  |  |  |
| RPL42A |  |  |  |
| CDC33 |  |  |  |
| RPS15 |  |  |  |
| RPS29B |  |  |  |

|  |
| --- |
| RPS5 |
| PSA1 |
| GDH1 |
| RDL1 |
| THS1 |
| ENO1 |
| RPL22A |
| RPS24A |
| RPL12A |
| RPS1A |
| CIT1 |
| RPL40B |
| SED1 |
| EFT2 |
| HIS4 |
| YJL133C-A |
| HHT2 |
| ADK1 |
| CCW12 |
| ECM33 |
| OLE1 |
| RPL38 |
| RPS20 |
| PMT2 |
| GIS2 |
| SSB2 |
| NUS1 |
| OPI3 |
| ARF2 |
| TSA1 |
| RPS0A |
| RPL34A |
| OCA1 |
| RPL35A |

